# The human reward system encodes the subjective value of ideas during creative thinking

**DOI:** 10.1101/2024.03.07.583926

**Authors:** Sarah Moreno-Rodriguez, Benoît Béranger, Emmanuelle Volle, Alizée Lopez-Persem

## Abstract

Creative thinking is composed of a generation phase, where individuals form candidate ideas, and an evaluation phase, where individuals monitor the quality of their ideas, in terms of both their originality and adequacy, to select the best one. Here, we conceptualize creative evaluation as a specific type of decision-making, where participants attribute subjective values to ideas to guide their choice. Yet, while subjective values and preferences have been the focus of many studies in classical decision-making, their involvement in creative decision-making remains largely unexplored. Combining creative generation tasks and rating tasks, the present study demonstrates that individuals assign subjective values to ideas and that these values depend on a relative balance of the ideas’ originality and adequacy, which is determined by individual preferences and predicts their creative abilities. Using functional Magnetic Resonance Imaging, we found that the human reward system encodes the subjective value of ideas, and that the Default Mode and the Executive Control Networks, rather than being split into idea generation and evaluation, respectively reflect the originality and adequacy of ideas. Interestingly, the relative functional connectivity of the Default Mode and Executive Control Networks with the human reward system correlates with the relative balance of adequacy and originality in individuals’ preferences. These results bridge a gap in the current literature by providing new evidence regarding the neural bases for originality and adequacy monitoring and add valuation to the incomplete behavioral and neural accounts of creativity, offering perspectives on the influence of individual preferences on creative abilities.

## 1. Introduction

From global issues to daily struggles, humans consistently face new challenges and must rely on their creativity to solve them. In neuroscience, creativity is defined as the ability to form ideas that are both original and adequate (Stein, 1953; F. Barron, 1955; Runco & Jaeger, 2012). Original ideas are infrequent, unusual, novel or unique. Adequate ideas are appropriate, efficient, and useful; they comprise the requirements to fulfill their purpose. Despite an objective definition of *what* is creative, *how* individuals achieve a creative production remains unclear (Green et al., 2023). Notably, two people will rarely come up with a similar solution to a problem nor prefer the same idea when asked for the best alternative (Benedek et al., 2016; Lloyd-Cox, Pickering, et al., 2022; Silvia, 2008). In fact, we still have a limited understanding of how individuals integrate originality and adequacy during creative performance and how individual preferences influence it.

The dominant neurocognitive view for creativity proposes a dual-process framework, decomposing the creative process into an idea-generation phase and an evaluation phase (Allen & Thomas, 2011; Barr et al., 2015; R. Beaty et al., 2014, 2016; Benedek & Jauk, 2018; Cassotti et al., 2016; Chrysikou, 2019; Ellamil et al., 2012; Kleinmintz et al., 2019; Sowden et al., 2015; Volle, 2018). In the generation phase, individuals form novel associations: they link seemingly unrelated concepts and produce a wide variety of potential solutions. This phase can be facilitated by memory structure (Benedek et al., 2017; Benedek & Fink, 2019; He et al., 2020; Ovando-Tellez et al., 2022), associative abilities (R. Beaty et al., 2014; R. E. Beaty, Zeitlen, et al., 2021; He et al., 2020; Wang et al., 2023) and cognitive flexibility (Kenett et al., 2018; Li et al., 2021), enabling individuals to navigate between different concepts during idea generation. In the evaluation phase, candidate ideas are assessed for originality and adequacy and selected according to the context. Evaluation processes are assumed to be deliberate and goal-directed and to rely on cognitive control mechanisms (Kleinmintz et al., 2019; Sowden et al., 2015; Wang et al., 2023). Interestingly, the evaluation of originality and adequacy might vary between individuals and the tasks at hand (Lloyd-Cox, Pickering, et al., 2022). However, the exact computations at play during the two phases, especially during the evaluation of ideas, remain undetermined.

In this study, we conceptualize the evaluation step of creativity as a specific case of decision-making: similar to food options in dietary choices, creative options are compared, and the most satisfactory one is selected (Lin & Vartanian, 2018; Lopez-Persem et al., 2023). We hypothesize that creative evaluation involves individual preferences and decision-making computations that methods from neuroeconomics can enable us to study. Specifically, we argue that creativity involves a valuation process.

In neuroeconomics, valuation consists of assigning a subjective value to an item - reflecting how valuable the item is, how much the subject likes it - and guides selection (Redish et al., 2016; Rushworth & Behrens, 2008). Similarly, we propose that the subjective value of a creative idea reflects the satisfaction it provides and plays a crucial role in guiding idea selection. In a previous work (Lopez-Persem et al., 2023), we demonstrated that creativity indeed involves valuation. In particular, that study first showed that the speed at which individuals produced ideas correlated to how much they liked them, suggesting that subjective valuation of ideas energizes creative production. Second, the value given to an idea was a function of its originality and adequacy, indicating that subjective valuation integrates the fundamental criteria of creativity. This integration varied across individuals, with some giving more weight to originality than adequacy. Finally, we showed that originality-inclined individuals yielded better creativity scores than the adequacy-inclined, showing the impact of valuation patterns on creative abilities. Overall, this seminal study empirically established that valuation plays a central role in creativity. In the current study, we replicate those results and aim to identify the neural underpinnings of the valuation processes involved in creative thinking.

At the neural level, creativity research has found the generation and evaluation phases to be associated with the Default Mode Network (DMN) and the Executive Control Network (ECN), respectively (R. Beaty et al., 2016; Ellamil et al., 2012; Kleinmintz et al., 2019). The DMN is a network composed of regions of the medial prefrontal cortex (mPFC), the posterior cingulate cortex (PCC), the precuneus, the inferior parietal lobule (IPL) and the temporal lobe. It is observed in individuals at rest, unfocused on their environment (Raichle, 2015). The DMN also has a role in essential cognitive functions, including associative thinking, daydreaming, mind-wandering, and self-referential thoughts (Zabelina & Andrews-Hanna, 2016). In the context of creativity, the DMN has proven to be central to idea generation (Liu et al., 2015; Lloyd-Cox, Chen, et al., 2022; Marron et al., 2018; Matheson et al., 2023; Mayseless et al., 2014, 2015; Wang et al., 2023). In parallel, creativity research has found the evaluation phase of the dual process to be associated with the ECN. The ECN includes regions of the dorsolateral prefrontal cortex (dlPFC), the anterior cingulate cortex (ACC), the inferior parietal lobule (IPL) and the temporal lobe. It has been consistently associated with cognitive control, working memory processes, goal-directed behavior, task switching and decision-making (Niendam et al., 2012). In creativity, research has shown that the ECN supports idea evaluation (Ellamil et al., 2012; Huang et al., 2018; Liu et al., 2015; Lloyd-Cox, Chen, et al., 2022; Matheson et al., 2023; Mayseless et al., 2014; Wang et al., 2023; Wu et al., 2022). Interestingly, the ECN and DMN typically exhibit mutually exclusive activity depending on whether the task is internally or externally focused (Anticevic et al., 2012). However, in the context of creativity, their cooperation has been widely observed and appears to be essential for creative thinking and performance (R. Beaty et al., 2016; Kleinmintz et al., 2019; R. Beaty et al., 2018; Kühn et al., 2014; R. Beaty et al., 2015).

Despite the extensive research on creative evaluation, it remains unclear where originality and adequacy are encoded during creative thinking and how this information is used to guide creative performance. Moreover, decision-making processes and their neural substrates have been overlooked, and a notable blind spot has emerged in our understanding of the neurocognitive bases of creative evaluation.

In neuroeconomics, research on the neural bases of decision-making processes shows that subjective values are encoded in the human reward system, also known as the brain valuation system (BVS) (Bartra et al., 2013; Lopez-Persem et al., 2020). The BVS overlaps the reward system identified in animal literature (Schultz, 2015). It comprises the ventromedial prefrontal cortex (vmPFC), the orbitofrontal cortex (OFC), and subcortical regions, particularly the ventral striatum (VS) and the ventral tegmental area (VTA). The valuation process in the BVS is both generic and automatic. Generic in the sense that the activity in the BVS encodes the value of an object regardless of its nature. Automatic in that the value is encoded in the BVS even when an individual is evaluating another attribute, such as the age of a painting (Lebreton et al., 2009; D. J. Levy & Glimcher, 2012; I. Levy et al., 2011; Lopez-Persem et al., 2020). Besides the BVS, the ECN is also engaged in decision-making processes: it ensures top-down control by adjusting the values of options based on the context of the decision (Hare et al., 2009). The ECN is also responsible for integrating the difference in values between options. This difference in values is referred to as the decision value, as it drives the selection process (Domenech et al., 2018).

Based on its generic properties and its cooperation with the ECN, we hypothesize that idea valuation during creativity is supported by the BVS. This is consistent with several findings in creativity research, demonstrating the involvement of BVS regions (Beversdorf, 2019; Huang et al., 2015; Takeuchi et al., 2010) or dopaminergic modulation (Aberg et al., 2016; Boot et al., 2017; Chermahini & Hommel, 2010; Zabelina et al., 2016) in creativity. Despite these results and the overall recognized importance of motivation in creativity, its neural bases and the probable involvement of the BVS in creativity have been largely overlooked.

In summary, we propose to investigate the neural underpinnings of idea evaluation and valuation during creativity. Using functional Magnetic Resonance Imaging (fMRI), we aimed to (i) demonstrate that the subjective value of an idea is encoded in the BVS and (ii) clarify where originality and adequacy are encoded in the brain during idea production.

To address these questions and hypotheses, we recruited forty participants who performed a creative word association task during an fMRI session. Participants also rated how likeable, original and adequate the associations were. We first replicated the behavioral results of our previous work (Lopez-Persem et al., 2023), showing that valuation plays a role in creativity. Then, we identified the brain regions underlying the valuation of ideas and the encoding of originality and adequacy during the rating tasks. Based on these functional localizers, we determined whether and where subjective values, originality and adequacy were encoded in the brain when producing creative ideas in the word association task.

## 1. Results

The study consisted of several successive tasks (Figure 1): a Free Generation of Associate Task (FGAT), a likeability rating task and a choice task, all performed in an MRI, followed by an originality and adequacy rating task and a battery of creativity tests completed outside of the MRI. Thirty-eight subjects were included in the analyses (19 females, age = 26.5 ±0.7 (M±SEM), years of education = 16.7 ±0.4 (M±SEM)).

**Figure 1:**
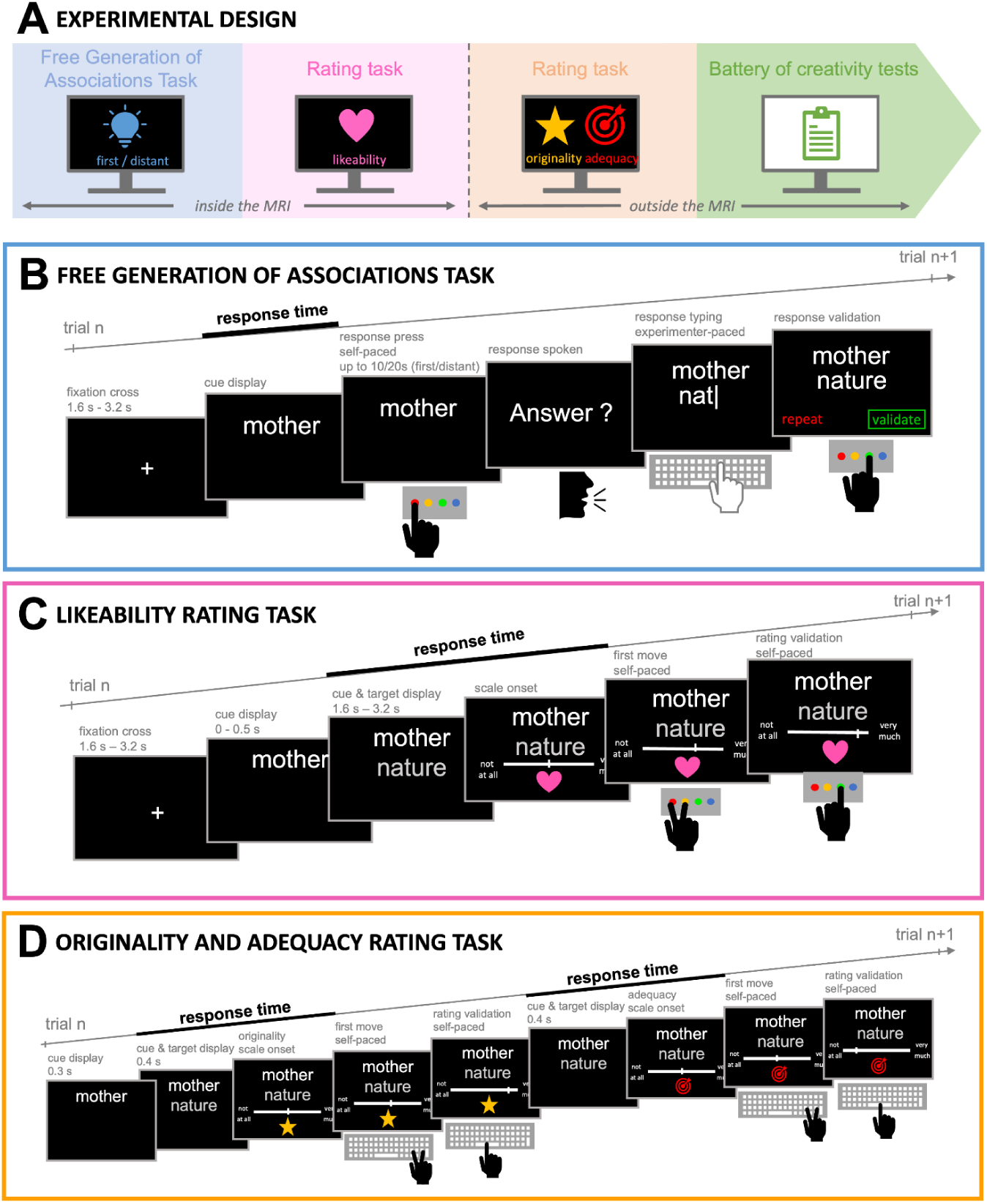
Experimental design. **(A)** Overview of the different tasks in chronological order **(B)** Free Generation of Association Task (FGAT). In two conditions (“first” and “distant”), participants had to provide, respectively: the first word that came to their mind when they saw the cue word or an unusual, original, but associated word. Participants in the MRI provided their responses orally to the experimenter, who typed them. They had the opportunity to repeat their response in case the experimenter had misheard. **(C)** Likeability rating task. Participants rated each association on a likeability scale from “not at all” to “very much” and validated the rating using MRI response buttons. **(D)** Originality and adequacy rating task. Participants rated each association on a scale going from “not at all” to “very much” using the keyboard arrows and validated the rating using the spacebar. In the MRI, all trials of all tasks began with a fixation cross, followed by the trial screen. Participants also completed a choice task in the MRI scanner, for which data were not analyzed for the current study.

### 2.1. Behavioral results

In the FGAT-first condition, we asked participants to give the first word that came to mind when reading a cue word. In the “distant” condition, they had to give a word that was farther from the cue word, resulting in more creative associations than in the “first” condition (Figure 1B). Then, participants rated how much they liked the associations (the association’s subjective value) and how original and adequate they found them on a scale of 0 to 100 (Figure 1C).

#### 2.1.1. Individuals achieve better adequacy and originality when trying to be creative

To characterize participants’ creative productions, we first explored the differences between FGAT “first” and “distant” responses, using the participants’ adequacy and originality ratings. As expected, “first” associations were highly adequate and little original (adequacy rating = 83.1±1.4; originality rating = 36.8±1.5, M±SEM across participants), while “distant” associations were highly adequate and original (mean adequacy rating = 73.1±1.4; mean originality rating = 63.2±1.3). In addition, we found that the difference in originality between “first” and distant” associations was greater than the difference in adequacy (paired two-tailed t-test: t(37)=10.4, p=2.10^-12^), indicating that “distant” associations were both original and adequate - i.e., creative - while “first” associations were mainly adequate (Figure 2A left).

**Figure 2:**
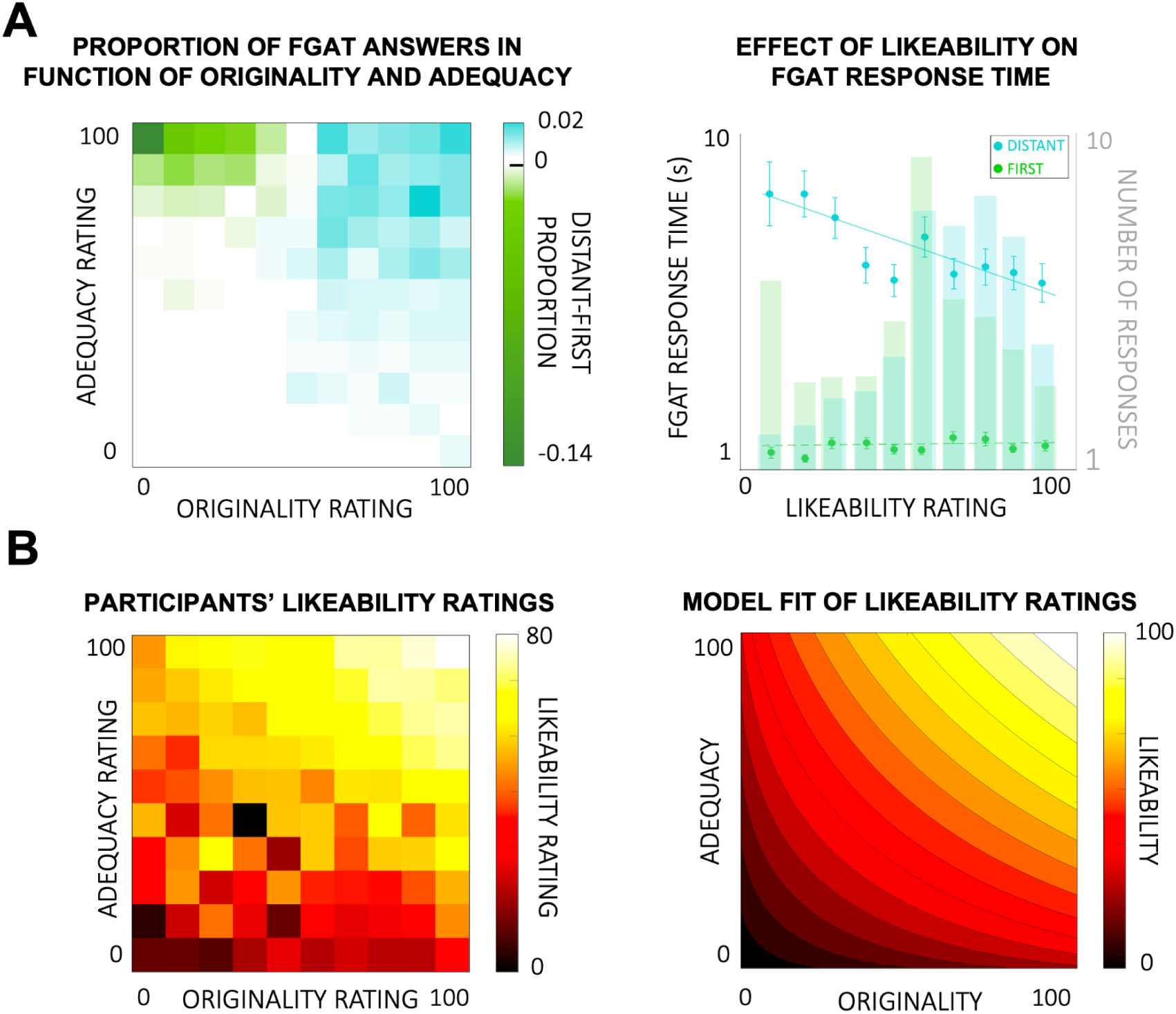
Behavioral results of creative idea production and evaluation. **(A)** Creative idea production. **Left:** Heatmap of the difference in proportion between “distant” and “first” associations per bin of adequacy and originality ratings. Bins with positive values count more “distant” answers than “first” answers and vice versa. **Right:** Correlation between response time and likeability ratings of the FGAT responses for the “first” and “distant” conditions. Background bars indicate the mean number of answers per bin of likeability ratings. Circles are bins of averaged participant data. Error bars are intersubject standard errors of the mean (SEM). Lines correspond to the linear regression fit at the group level in the “distant” condition (significant fit, p<0.05) and the “first” condition (non-significant fit, p>0.05). (**B)** Creative idea evaluation. **Left:** Heatmap of participants’ average likeability ratings per bin of adequacy and originality ratings. **Right**: CES model fit of likeability ratings in function of adequacy and originality ratings. The verticality of isolines is captured by the α parameter (preference for originality), while the convexity of isolines is captured by the δ parameter (preference for a trade-off of both dimensions rather than extremes).

Then, using an objective measure - i.e., the associative frequency of responses, measured with the dictionary of verbal associations “Dictaverf”, we also found that “distant” responses were less frequent than “first” responses (paired two-tailed t-test: t(37)=-23.7, p<.001), confirming that the “distant” condition elicited more remote, creative ideas.

#### 2.1.2. Individuals provide preferred responses faster

Then, we looked for a behavioral signature of preferences during idea production. We inspected response times (RT) in the FGAT, i.e., the thinking time between when the participant saw the cue word and when they pressed the button to give their response (Figure 1B).

We found that participants generated the ideas they liked faster than the ones they disliked in the “distant” condition (RT_distant_ regressed against likeability: β_distant_=-0.18±0.02, one-sample two-tailed t-test: t(37)=-7.48, p=7.10^-9^) but not in the “first” condition (RT_first_ regressed against likeability: β_first_=0.04±0.04, one-sample two-tailed t-test: t(37)=1.26, p=0.2). The effect of likeability on response times was significantly different between conditions (effect of likeability on “distant” versus “first” RT - i.e. β_distant_-β_first_: paired two-tailed t-test: t(37)=-4.74, p=3.10^-5^, Figure 2A right). Note that this result holds when adding an additional regressor of squared standardized likeability as a proxy for confidence (H. Barron et al., 2015; Lebreton et al., 2015) (Supplementary Results).

Overall, the effect of likeability on the FGAT-distant response times suggests an implicit involvement of valuation during creative idea production. When generating ideas, participants intrinsically value them, which is reflected in their response time.

#### 2.1.3. Individuals prefer responses that are both adequate and original

Next, we investigated how likeability judgements integrated adequacy and originality judgements. We observed that likeability increased with adequacy and originality: the more original and adequate an association was, the more it pleased participants (Figure 2B left).

We used the Constant Elasticity of Substitution (CES) model (Andreoni & Miller, 2002; Lopez-Persem et al., 2017), which had provided the best fit in our previous study using the same experimental design (Lopez-Persem et al., 2023). This model (see *Material and Methods 4.3.3* for the model equation) has two free parameters: alpha (α), which captures the weight given to originality relative to adequacy in the likeability ratings, and delta (δ), the preference for an equilibrium of originality and adequacy. We confirmed that the CES fitted the present dataset (mean fit ± standard error of the mean (SEM): r^2^=0.42±0.03; one-sample two-tailed t-test: t(37)=15.43, p<0.001; Figure 2B right).

For each participant, we estimated the model’s free parameters (hereafter referred to as valuation parameters). At the group level, we found that participants’ likeability ratings resulted from a balanced trade-off of originality and adequacy (no preference for originality: *α* = 0.46±0.04, one-sample two-tailed t-test against 0.5: t(37)=-0.92, p=0.36; preference for equilibrium: *δ* = 0.44±0.21, one sample two-tailed t-test against 1: t(37)=-2.67, p=0.011). In other words, associations that were both original and adequate (rather than very original or very adequate) yielded higher likeability ratings at the group level. This suggests that valuation similarly and conjointly considers an idea’s originality and adequacy, which is also consistent with the formal definition of creativity.

As a confirmatory analysis, we checked the relationship between the α parameter and the likeability of rare responses. Initially, we regressed likeability ratings against response frequency but found no significant effect at the group level (β_likeability-frequency_=-0.01±0.03, one-sample two-tailed t-test: t(37)=-0.4, p=0.7; Figure S1A). However, post-hoc analyses on individual estimations of the α parameter revealed a high interindividual variability (coefficient of variation in *α* = 58%). Interestingly, this was linked to a strong negative correlation between the α parameter and the β_likeability-frequency_ (r=-0.82, p=1.10^-10^, Figure S1B). This correlation means that participants who favored originality also preferred rare associations, which nuances the null relationship between likeability and frequency at the group level and is consistent with our interpretation of the α parameter.

#### 2.1.4. Individuals’ preferences are related to creative abilities

Using a battery of tests and questionnaires, we estimated participants’ creative abilities and investigated how they related to the valuation parameters of the CES using canonical correlation analysis.

We found one canonical variable showing significant dependence between the set of valuation parameters and the set of creative ability measures (r=0.52, p=0.025, Figure 3A). Both valuation parameters significantly contributed to the canonical variable (*α*: r=0.34, p=0.04; and *δ*: r=-0.50, p=0.0013, Figure 3B) and among the creativity set, all but one of the creativity scores contributed to the canonical variable (associative fluency task score: r=0.34, p=0.035; CAT score: r=0.21, p=0.20; drawing task score: r=0.44, p=0.006; ICAA score: r=0.38, p=0.018, Figure 3C).

**Figure 3:**
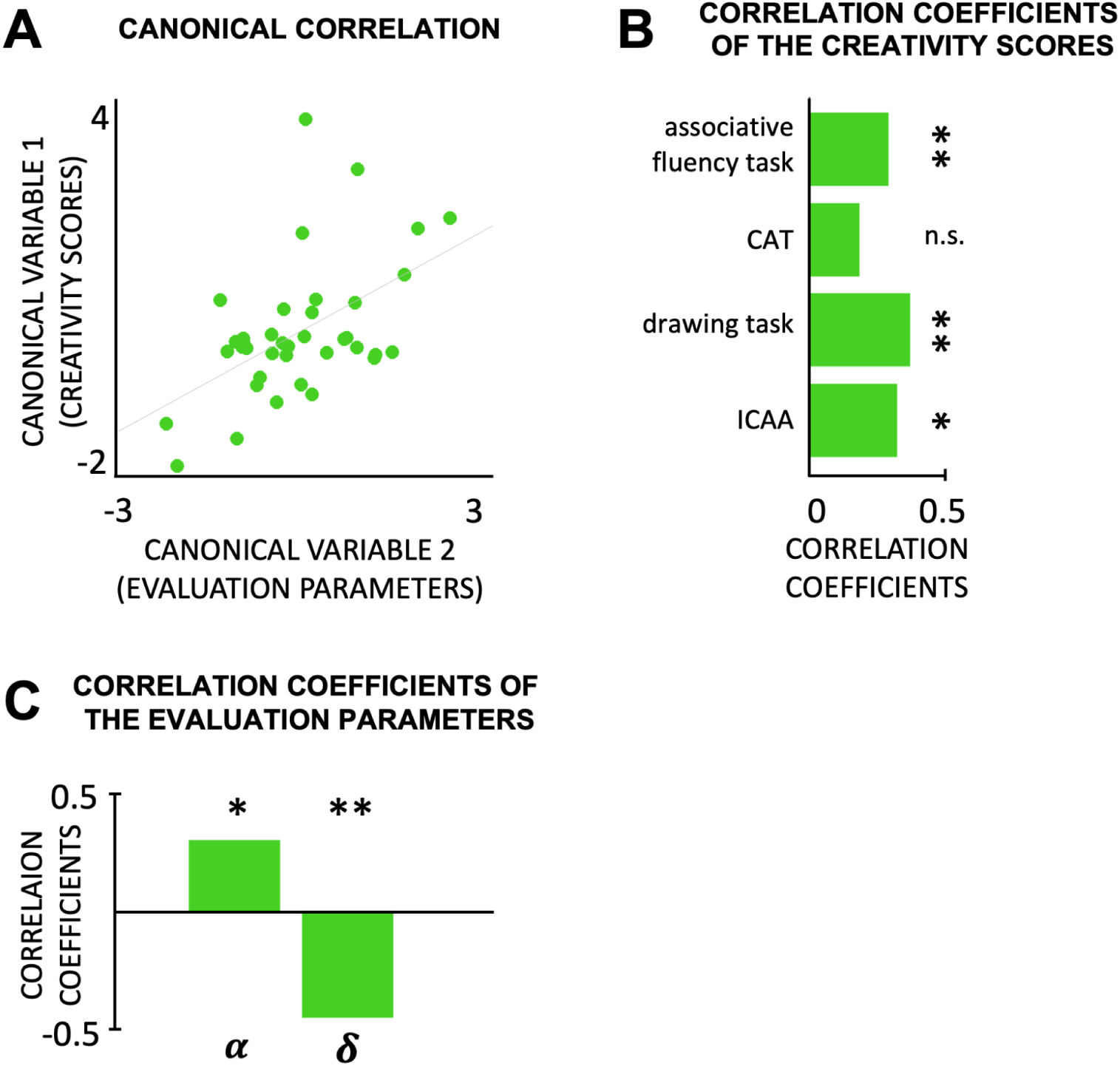
Canonical correlation between evaluation parameters and creative abilities. **(A)** Canonical correlation between creativity scores in battery tests and evaluation parameters (*α* and *δ*) from the behavioral analyses. Each circle represents one participant. The line represents a significant correlation (p<0.05). (**B)** Correlation coefficients of the creativity scores from the battery of tests. (**C)** Correlation coefficients of the evaluation parameters. * and ** indicate statistical significance (respectively p<0.05 and p<0.01). n.s. stands for not significant (p>0.05).

These results indicate that individuals’ valuation pattern is related to their creative abilities: participants who valued originality (i.e., had a high α parameter) and its balance with adequacy (i.e., a low δ parameter) yielded higher scores in the battery of creativity tests.

### 2.2. Neuroimaging results

Behavioral results indicate that the production of creative associations involves participants’ preferences. We then wanted to explore whether and which neural regions encoded subjective values (likeability) during idea production. Additionally, given that likeability relies on the adequacy and originality of ideas, we also explored the neural representations of those two dimensions. We first examined the relationship between each variable and neural activity during the likeability rating task to define functional localizers. Then, we investigated whether the identified brain regions were also involved in encoding those variables during creative idea production in the FGAT-distant condition.

#### 2.2.1. Likeability ratings are encoded in the BVS during idea evaluation and production Defining the functional localizer of likeability evaluation

We used a whole-brain parametric modulation approach to investigate the neural representation of likeability during the likeability rating task, with likeability ratings as regressors. The GLM we tested contains the likeability of the responses (from this likeability rating task) as a parametric modulator of a categorical boxcar regressor between the onset of the cue (i.e., the association) and the first button press to provide the likeability rating. We found a significant correlation between the associations’ likeability ratings and the blood-oxygen-level-dependent (BOLD) signal in regions classically identified as parts of the BVS (Figure 4A): the vmPFC (peak Montreal Neurological Institute (MNI) coordinates: [x=-8 y=48 z=-10], t(37)=8.20, pFWE=5.10^-15^) and the ventral striatum of the left ([-14, 22, 0], t(37)=7.55, pFWE=1.10^-7^) and right hemisphere ([12, 22, -2], t(37)=6.98, pFWE=4.10^-6^, see Table 1 for all significant clusters). This aligns with our hypothesis that likeability ratings are encoded in the BVS during idea valuation, similarly to any other item valuation.

**Figure 4:**
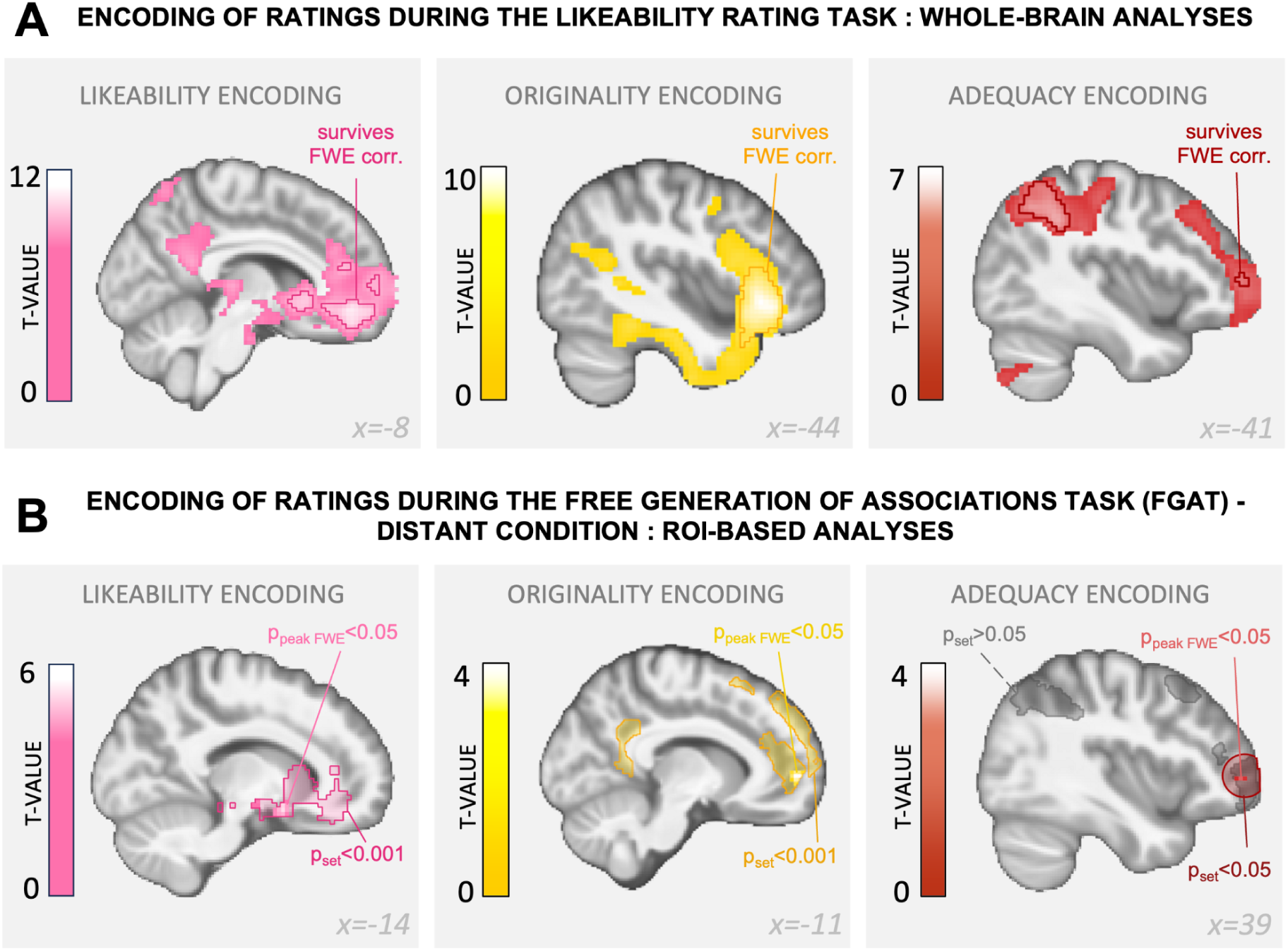
Neuroimaging results of likeability, originality and adequacy encoding during the likeability rating task and the FGAT-distant. **(A)** Neural encoding of likeability ratings (pink), originality ratings (yellow) and adequacy ratings (red) during the likeability rating task. The color code indicates the T-value of one-sample t-tests, p<0.001 uncorrected. Circled regions survived cluster-level FWE correction (p<0.05). See Table 1 for all significant clusters. **(B)** Neural encoding of likeability ratings (pink), originality ratings (yellow) and adequacy ratings (red) during the FGAT-distant. Circled regions are the overlap of the regions in A and their most similar atlas network (as depicted in Figure 5). They were used as inclusive masks for small volume correction, to identify significant peaks with FWE correction (p<0.05). Colored circled regions are significant at the set level (p<0.001); grey circled regions are not significant at the set level (p>0.05). The color code of the peaks within the masks indicates the T-value of one-sample t-tests, displayed at p<0.001 but surviving FWE correction (p<0.05) within the mask. See Table 2 for more information.

**Table 1:** All significant clusters in the SPMs of the parametric modulation analyses during the rating task. The clusters listed here survived whole-brain cluster-level FWE correction (p<0.05). Coordinates x, y and z refer to the Montreal Neurological Institute space. The last column indicates whether the peak activation location falls within a network of interest. DMN and ECN are defined by Yeo et al.’s atlas (2011), and BVS is defined by the term-based meta-analyses of the Neurosynth platform https://neurosynth.org/analyses/terms/reward/). Only the results in bold are reported in the text.

| parametric modulator: likeability rating |  |  |  |  |  |  |  |  |  |
| --- | --- | --- | --- | --- | --- | --- | --- | --- | --- |
| cluster label | cluster size | peak side | x | y | z | Z-score | T-value | cluster p-value | network of interest? |
| occipital cortex | 804 | R | 6 | -82 | -2 | >8 | 12.81 | <10<sup>-16</sup> | none |
| **ventromedial prefrontal cortex** | **324** | **L** | **-8** | **48** | **-10** | **6.15** | **8.20** | **5.10-15** | **DMN** |
| **ventral striatum** | **91** | **L** | **-14** | **22** | **0** | **5.84** | **7.55** | **1.10-7** | **BVS** |
| **ventral striatum** | **57** | **R** | **12** | **22** | **-2** | **5.54** | **6.98** | **4.10-6** | **BVS** |
| cerebellum | 30 | R | 12 | -52 | -50 | 5.36 | 6.65 | 1.10-4 | none |
| anterior cingulate cortex | 18 | L | -11 | 42 | 16 | 5.33 | 6.60 | 6.10-4 | DMN |
| medial frontal pole | 13 | L | -8 | 60 | 6 | 4.94 | 5.92 | 0.002 | DMN |
| midbrain | 1 | L | -6 | -18 | -24 | 4.89 | 5.85 | 0.027 | BVS |
| parametric modulator: originality rating |  |  |  |  |  |  |  |  |  |
| cluster label | cluster size | peak side | x | y | z | Z-score | T-value | cluster p-value | network of interest? |
| **inferior frontal gyrus** | **1163** | **L** | **-44** | **28** | **-4** | **7.44** | **11.46** | **<10<sup>-16</sup>** | **none** |
| **middle temporal gyrus** | **543** | **L** | **-56** | **-15** | **-12** | **6.64** | **9.30** | **<10<sup>-16</sup>** | **DMN** |
| **dorsomedial prefrontal cortex** | **320** | **L** | **-6** | **48** | **16** | **6.55** | **9.10** | **3.10-15** | **DMN** |
| **lateral orbitofrontal cortex** | **105** | **R** | **39** | **30** | **-10** | **5.85** | **7.57** | **3.10-8** | **BVS** |
| **pre-supplementary motor area** | **72** | **L** | **-4** | **20** | **58** | **5.80** | **7.48** | **6.10-7** | **DMN** |
| ventral tegmental area | 27 | L | -4 | -22 | -20 | 5.78 | 7.43 | 1.10-4 | BVS |
| lateral orbitofrontal cortex | 35 | R | 26 | 18 | -22 | 5.63 | 7.16 | 5.10-5 | none |
| medial frontal pole | 31 | L | -4 | 62 | -14 | 5.54 | 6.99 | 8.10-5 | BVS |
| occipital cortex | 49 | L | -18 | -95 | -7 | 5.54 | 6.99 | 8.10-6 | none |
| cerebellum | 31 | R | 2 | -55 | -42 | 5.49 | 6.89 | 8.10-5 | none |
| brainstem | 150 | L | -4 | -30 | -2 | 5.47 | 6.84 | 5.10-10 | BVS |
| occipital cortex | 87 | R | 24 | -90 | -4 | 5.38 | 6.68 | 1.10-7 | none |
| cerebellum | 39 | R | 19 | -78 | -40 | 5.37 | 6.66 | 3.10-5 | none |
| fusiform gyrus | 10 | L | -34 | -8 | -42 | 5.15 | 6.29 | 0.002 | none |
| amygdala | 7 | L | -16 | -2 | -17 | 4.95 | 5.95 | 0.005 | BVS |
| temporal pole | 3 | R | 42 | 15 | -27 | 4.91 | 5.88 | 0.013 | none |
| fusiform gyrus | 2 | L | -38 | -10 | -37 | 4.85 | 5.78 | 0.018 | none |
| posterior cingulate cortex | 3 | L | -8 | -50 | 30 | 4.82 | 5.74 | 0.013 | DMN |
| globus pallidus | 3 | R | 14 | 0 | -14 | 4.82 | 5.73 | 0.013 | BVS |
| inferior frontal gyrus | 1 | L | -48 | 18 | 20 | 4.77 | 5.65 | 0.026 | ECN |
| cerebellum | 1 | R | 22 | -40 | -47 | 4.75 | 5.63 | 0.026 | none |
| parametric modulator: adequacy rating |  |  |  |  |  |  |  |  |  |
| cluster label | cluster size | peak side | x | y | z | Z-score | T-value | cluster p-value | network of interest? |
| **precuneus** | **192** | **L** | **-6** | **-70** | **38** | **5.96** | **7.80** | **3.10-11** | **ECN** |
| **intraparietal lobule** | **398** | **L** | **-41** | **-48** | **38** | **5.95** | **7.77** | **1.10-16** | **ECN** |
| **posterior cingulate cortex** | **129** | **L** | **-6** | **-25** | **30** | **5.86** | **7.59** | **5.10-9** | **ECN & BVS** |
| **lateral frontal pole** | **110** | **L** | **-36** | **52** | **3** | **5.74** | **7.36** | **4.10-6** | **ECN** |
| **intraparietal lobule** | **122** | **R** | **44** | **-38** | **43** | **5.57** | **7.04** | **9.10-9** | **ECN** |
| **inferior temporal sulcus** | **34** | **L** | **-61** | **-30** | **-17** | **5.28** | **6.50** | **7.10-5** | **DMN** |
| cerebellum | 15 | R | 44 | -65 | -50 | 5.23 | 6.41 | 0.001 | none |
| **lateral frontal pole** | **55** | **R** | **34** | **62** | **6** | **5.12** | **6.24** | **5.10-6** | **ECN** |
| lingual gyrus | 10 | R | 6 | -78 | -7 | 5.08 | 6.17 | 0.003 | none |
| lingual gyrus | 10 | R | 16 | -72 | -12 | 5.04 | 6.10 | 0.003 | none |
| medial frontal pole | 9 | L | -8 | 75 | 3 | 4.94 | 5.93 | 0.003 | none |
| supramarginal gyrus | 2 | L | -51 | -35 | 43 | 4.78 | 5.66 | 0.019 | ECN |
| lateral frontal pole | 2 | R | 42 | 62 | -12 | 4.76 | 5.64 | 0.019 | none |

**Table 2:** All clusters in the SPMs of the parametric modulation analyses during the FGAT-distant. Only the results in bold are reported in the text. p-values are FWE corrected. Coordinates x, y and z refer to the Montreal Neurological Institute space. Only the results in bold are reported in the text.

| parametric modulator: **likeability rating** |  |  |  |  |  |  |  |  |  |
| --- | --- | --- | --- | --- | --- | --- | --- | --- | --- |
| mask |  |  |  |  |  |  | set p-value |  |  |
| overlap of the likeability rating localizer with the BVS (Neurosynth metanalysis) |  |  |  |  |  |  | **0.007** |  |  |
| cluster label | cluster size | peak side | x | y | z | Z-score | T-value | peak p-value | cluster p-value |
| ventral striatum | 29 | L | -14 | 8 | -14 | 04.09 | 4.64 | 0.015 | **0.0501** |
| parametric modulator: **originality rating** |  |  |  |  |  |  |  |  |  |
| mask |  |  |  |  |  |  | set p-value |  |  |
| overlap of the originality rating localizer with the DMN (Yeo et al., 2011) excluding the BVS |  |  |  |  |  |  | **3.10-4** |  |  |
| cluster label | cluster size | peak side | x | y | z | Z-score | T-value | peak p-value | cluster p-value |
| medial frontal pole | 20 | L | -11 | 55 | 0 | 4.1 | 4.68 | 0.044 | **0.215** |
| parametric modulator: **adequacy rating** |  |  |  |  |  |  |  |  |  |
| mask |  |  |  |  |  |  | set p-value |  |  |
| overlap of the adequacy rating localizer and the ECN (Yeo et al., 2011) excluding the BVS |  |  |  |  |  |  | **0.292** |  |  |
| cluster label | cluster size | peak side | x | y | z | Z-score | T-value | peak p-value | cluster p-value |
| dorsolateral prefrontal cortex | 8 | R | 44 | 42 | 13 | 3.77 | 4.20 | 0.127 | 0.355 |
| lateral frontal pole | 8 | R | 42 | 55 | 0 | 3.30 | 3.59 | 0.445 | 0.355 |
| mask |  |  |  |  |  |  |  |  |  |
| 10-voxel radius sphere centered on the 1<sup>st</sup> peak activation in the adequacy rating localizer (x=-6 y=-70 z=38) |  |  |  |  |  |  |  | no suprathreshold clusters |  |
| mask |  |  |  |  |  |  |  |  |  |
| 10-voxel radius sphere centered on the 2<sup>nd</sup> peak activation in the adequacy rating localizer (x=-41 y=-48 z=38) |  |  |  |  |  |  |  | no suprathreshold clusters |  |
| mask |  |  |  |  |  |  |  |  |  |
| 10-voxel radius sphere centered on the 3<sup>rd</sup> peak activation in the adequacy rating localizer (x=-36 y=52 z=3) |  |  |  |  |  |  |  | no suprathreshold clusters |  |
| mask |  |  |  |  |  |  |  |  |  |
| 10-voxel radius sphere centered on the 4<sup>th</sup> peak activation in the adequacy rating localizer (x=44 y=-38 z=43) |  |  |  |  |  |  |  | no suprathreshold clusters |  |
| mask |  |  |  |  |  |  | set p-value |  |  |
| 10-voxel radius sphere centered on the 5<sup>th</sup> peak activation in the adequacy rating localizer (x=34 y=62 z=6) |  |  |  |  |  |  | **0.049** |  |  |
| cluster label | cluster size | peak side | x | y | z | Z-score | T-value | peak p-value | cluster p-value |
| lateral frontal pole | 2 | R | 39 | 58 | 0 | 3.13 | 3.37 | **0.045** | 0.035 |

##### Comparing the functional localizer of likeability evaluation to atlas functional networks

To confirm that likeability evaluation involved the BVS, we compared our localizer with the BVS (as defined by the term-based meta-analyses of the Neurosynth platform https://neurosynth.org/analyses/terms/reward/), as well as 7 intrinsic functional networks from the Yeo atlas (2011). For this comparison, we quantified the number of voxels in common between our localizer and each atlas network (Figure 5). We found that the regions reflecting likeability ratings mostly overlapped with the BVS network (31 688 voxels out of 112 370, i.e. 28% of overlap), more so than with the other 7 networks, which aligned with our observations and hypotheses. For the following analyses, we defined the BVS region of interest (ROI) as the overlapping voxels of the functional localizer and the BVS Neurosynth network (the dark pink areas in Figure 5).

**Figure 5:**
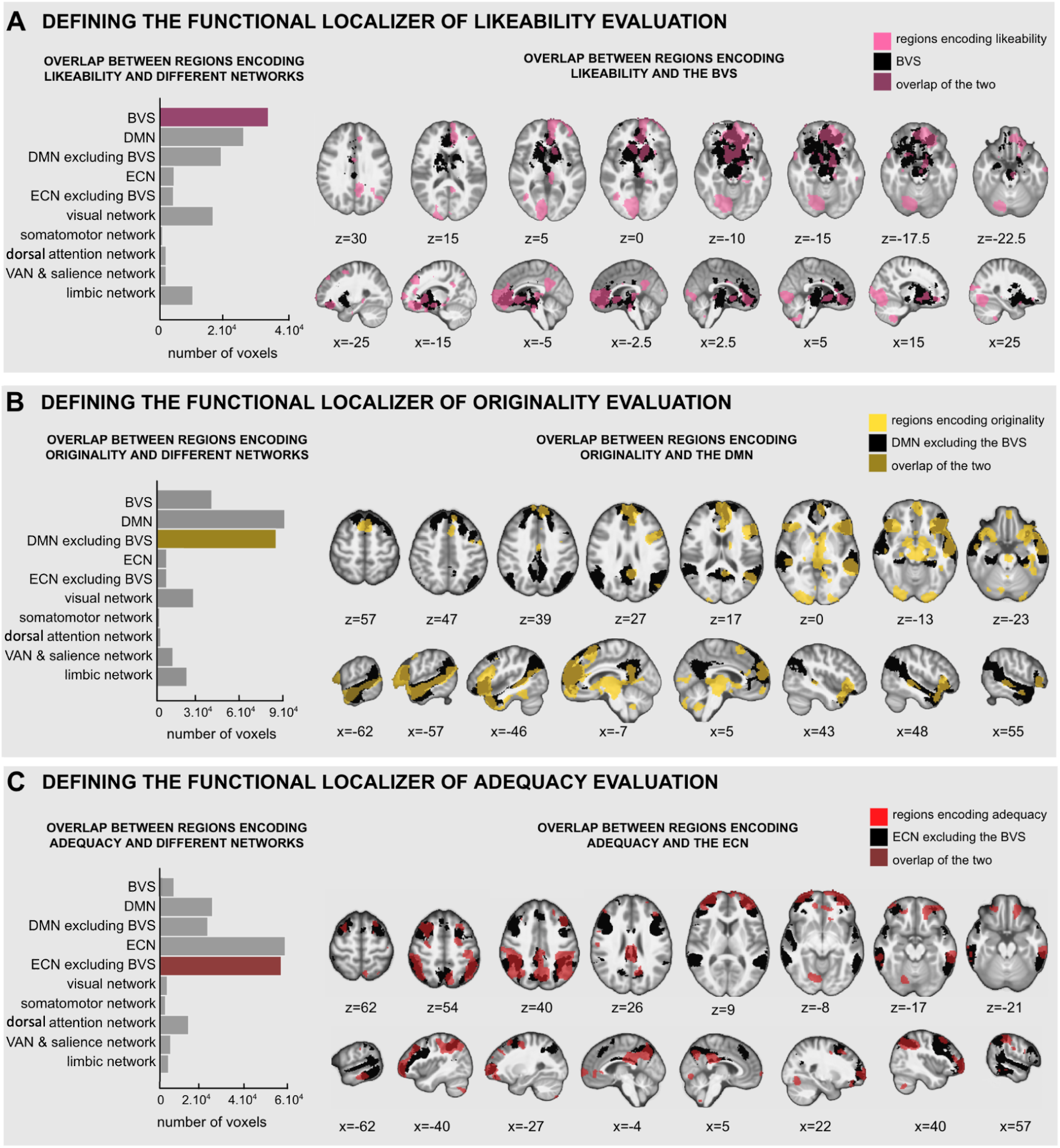
Defining the function localizers of likeability, originality and adequacy evaluation using brain networks overlaps Left: Number of common voxels between the statistical maps of the parametric modulation of the likeability rating task by **(A)** likeability, **(B)** originality and **(C)** adequacy ratings, with networks of interest as defined by Yeo et al. (2011) and Neurosynth. See Table S1 for all inter-network overlaps. **Right:** Visual of the overlap between the regions encoding each dimension (color) and the principal network (black) as defined by Yeo et al. (2011) and Neurosynth.

##### Locating the neural encoding of likeability during idea production

We performed ROI-based parametric modulation analyses on the FGAT-distant to determine whether likeability was also encoded in the BVS during the production of creative ideas and specify which regions in particular. The GLM we tested contains the likeability of the responses (obtained in the rating task) as a parametric modulator of a categorical boxcar regressor between the onset of the cue and the button press to provide the creative response. We found a significant correlation with likeability within the BVS ROI (set p-value=0.007; see Table 2 for details). Plus, when investigating the clusters within the ROI, we found a significant correlation between the likeability of the response and the BOLD signal in the ventral striatum ([-14, 8, -14], t(37)=4.64, peak p-value (FWE)=0.015; Figure 4B), which confirms our hypothesis that valuation is at play during idea production via the encoding of likeability in the BVS. Note that this result was controlled for response times (Supplementary Results and Table S2).

Interestingly, the same GLM did not yield any significant results when applied to the FGAT-first condition (peak p-values (FWE)>0.5), regardless of whether we used the ROI as an inclusive mask or not. This suggests that valuation was only at play during the creative idea production of the FGAT-distant condition, as opposed to when subjects formed spontaneous associations during the FGAT-first condition.

Together, these analyses confirm our central hypothesis that the evaluation phase of creativity partly relies on the assignment of subjective values encoded in the BVS. In the following sections, we investigated how originality and adequacy of ideas were represented in the brain.

#### 2.2.2. Originality and adequacy ratings are encoded in the DMN and the ECN during idea evaluation and production

##### Defining the functional localizer of originality and adequacy evaluation

To investigate the neural representation of adequacy and originality of ideas, we followed the same parametric modulation approach as before, this time with originality and adequacy ratings as parametric modulators. As participants rated originality and adequacy outside the scanner, we used the likeability rating task to create localizers of adequacy and originality judgements.

We found a significant correlation between the associations’ originality ratings and the BOLD signal in regions classically identified as parts of the DMN during the likeability rating task (Figure 4A): the inferior frontal gyrus ([-44, 28, -4], t(37)=11.46, pFWE<10^-16^), the middle temporal gyrus ([-56, -15, -12], t(37)=9.30, pFWE<10^-16^), the dorsomedial prefrontal cortex ([-6, 48, 16], t(37)=9.10, pFWE=3.10^-15^), the lateral orbitofrontal cortex ([39, 30, -10], t(37)=7.57, pFWE=3.10^-8^), and the pre-supplementary motor area ([-4, 20, 58], t(37)=7.48, pFWE=6.10^-7^).

In parallel, we found a significant correlation between the associations’ adequacy ratings and the BOLD signal in regions classically identified as parts of the ECN during the likeability rating task (Figure 4A): the precuneus ([-6, -70, 38], t(37)=7.80, pFWE=3.10^-11^), the left and right intraparietal lobule ([-41, -48, 38], t(37)=7.77, pFWE=1.10^-16^; [44, -38, 43], t(37)=7.04, pFWE=9.10^-9^), the posterior cingulate cortex ([-6, -25, 30], t(37)=7.59, pFWE=5.10^-9^), the left and right lateral frontal pole ([-36, 52, 3], t(37)=7.36, pFWE=4.10^-6^; [34, 62, 6], t(37)=6.24, pFWE=5.10^-6^), and the inferior temporal sulcus ([-61, -30, -17], t(37)=6.50, pFWE=7.10^-5^ - see Table 1 for all significant clusters). This aligns with our hypothesis that originality and adequacy ratings are encoded during idea evaluation, although we had no specific hypotheses regarding the respective underlying brain networks.

##### Comparing the functional localizer of originality and adequacy evaluation to atlas functional networks

To position the identified regions in brain networks and define ROI masks, we followed the same approach as for the likeability localizer (Figure 5). We found that the regions reflecting originality ratings mostly overlapped with the DMN (82 262 voxels out of 248 106, i.e. 33% of overlap) and that the regions reflecting adequacy ratings mostly overlapped with the ECN 59 921 voxels out of 172 662, i.e. 35% of overlap), more so than with the other 7 networks, which confirmed our visual observations (Figure 5 and Table S1 for all overlaps). For the following analyses, we defined the DMN and ECN ROIs as the overlapping voxels between the functional localizer and its corresponding atlas functional network while excluding the voxels that were in common with the BVS ROI (the dark yellow and dark red areas in Figure 5). Figure 5 also mentions “ECN excluding BVS” and “DMN excluding BVS” as we removed the BVS regions overlapping with Yeo et al.’s (2011) ECN and DMN from their respective ROIs in the following analyses to improve specificity.

##### Locating the neural encoding of originality and adequacy ratings during idea production

To investigate whether the DMN and ECN respectively represented originality and adequacy during the production of creative ideas, we followed the same approach as for likeability. We performed ROI-based parametric modulation analyses during the FGAT-distant to determine whether each network encoded its associated dimension during the production of creative ideas and specify which regions in particular.

We found a significant correlation with originality within the DMN ROI (set p-value=3.10^-4^; see Table 2 for details). Plus, when investigating the clusters within the ROI, we found a significant correlation between the originality of the response and the BOLD signal in the medial frontal pole ([-11, 55, 0], t(37)=4.68, peak p-value(FWE)=0.044; Figure 4B).

For adequacy, the set p-value was not significant (p=0.29), and we found no significant correlation between the associations’ adequacy and the BOLD signal anywhere at the cluster level. We resorted to using small-volume correction with a sphere centered on the five activation peaks from the localizer. We found a significant correlation between the associations’ adequacy and the BOLD signal only in the sphere located in the lateral frontal pole (sphere center coordinates: [34, 62, 6], peak activation coordinates: [39, 58, 0], t(37)=3.37, peak pFWE=0.045, set p-value=0.049; Figure 4B; see Table 2 for details).

Overall, these analyses first revealed that the DMN and the ECN respectively encoded originality and adequacy values during the likeability rating task, i.e. even when we had not instructed the participants to consider these two dimensions. In other words, participants tracked the association’s originality and adequacy while rating how much they liked it. Second, this encoding in the DMN and ECN was found again during the creativity task, suggesting that creative production also involves an evaluation of originality and adequacy on top of likeability.

#### 2.2.3. The contribution of the DMN and ECN to the BVS activity mirrors the contribution of originality and adequacy to the likeability ratings

Building on the encoding of originality and adequacy by the DMN and ECN, we tested whether the contribution of ECN and DMN activities to the BVS activity followed the same relationship as the contribution of adequacy and originality to likeability judgements (Figure S3). For each participant, we extracted the timeseries of activity for each network and fitted the CES value function to the fMRI data, with the BVS timeseries recorded during the FGAT-distant condition as the dependent variable and DMN and ECN timeseries as the independent variables (Figure S3A). This allowed us to estimate a neural *α* parameter and a neural *δ* parameter. Across individuals, we found a significant correlation between the *α*_neural_ and *α*_behavioural_ parameters (r=0.39, p=0.023) but not between the *δ* parameters (r=0.02, p=0.9) (Figure S3B). Interestingly, when performing the same analysis with the timeseries extracted from the resting-state session conducted at the end of the experiment, we still found a significant correlation between the *α*_neural_ and *α*_behavior_ parameters (r=0.46, p=0.0049) and again, no significant correlation for *δ* (r=-0.11, p=0.52) (Figure S3C). Together, these results suggest that the pattern of functional connectivity between the BVS, the ECN and the DMN reflects individual preferences, both during the creativity task and at rest.

## 2. Discussion

### Summary of findings

In this study, we used a creative word association task and several rating tasks to reveal the implication and the neural substrates of subjective valuation during the production of creative ideas. Our behavioral results replicate our prior work by showing that the subjective value of an idea depends on its originality and adequacy. We also replicate that valuation varies across individuals, resulting in the identification of individual patterns that relate to creative abilities. At the neural level, we show that i) subjective valuation is supported by the BVS during creative idea production; ii) originality and adequacy evaluations are respectively represented in the DMN and the ECN during creative idea production and iii) the DMN and ECN contribute to the BVS activity in a fashion that mirrors behavioral preferences and supports individual variability.

These results reveal the underlying neurocognitive mechanisms of the evaluation step of creativity and validate our hypothesis that individual preferences play a role in creativity.

### The role of the BVS in valuation and decision-making

We showed that the subjective value of an idea is represented in individuals’ BVS when we explicitly asked them to rate how much they liked an idea, but also when they focused on producing a creative idea.

This central result is consistent with two BVS properties reported in decision-making literature (Lopez-Persem et al., 2020). First, the involvement of the BVS in the valuation of creative ideas is consistent with its generic property: some have demonstrated that the BVS could reflect the subjective value of different kinds of items, no matter their nature (Lebreton et al., 2009; Motoki et al., 2019). This property aligns with the neural common currency framework, which states that the values of different and disparate types of items can be represented in the same brain regions and allows for their comparison (D. J. Levy & Glimcher, 2012). Second, the valuation of ideas in the BVS during creative production is consistent with the automatic property of the network: the BVS can reflect an item’s value even when an individual is not explicitly instructed to rate how much they like it. Even when an individual is focused on another task, for instance, when passively viewing a stimulus (I. Levy et al., 2011), when determining a person’s age (Lebreton et al., 2009; Lopez-Persem et al., 2020) - or in our case, when producing a creative word association - the BVS still reflects how much individuals like the item they are processing.

In parallel, the role of the BVS - and, in particular, the ventromedial prefrontal cortex (vmPFC) - has been challenged by recent studies. They argue that rather than representing one’s subjective value for the object they are processing (Bartra et al., 2013; Kable & Glimcher, 2009; Lebreton et al., 2009; Lopez-Persem et al., 2020; Rangel & Hare, 2010), the BVS could be representing slightly different variables, such as one’s goal value (Castegnetti et al., 2021; Daniel & Pollmann, 2014; De Martino & Cortese, 2023; Holton et al., 2023; Juechems & Summerfield, 2019; Park et al., 2021; Trudel et al., 2021) or one’s confidence in the rating (Clairis & Pessiglione, 2022; Lee et al., 2023; Rouault et al., 2022). In the current study, control analyses found no significant evidence indicating a confound with confidence. On the other hand, our study design does not allow us to differentiate between subjective and goal values. Subjective value differs from goal value in that the former is generally stable, regardless of the context, while the latter highly depends on one’s current goal. For example, the subjective value of a boat might always be minimal for someone not interested in traveling on the sea, but its goal value would increase dramatically if that person were stranded on a desert island.

This distinction might be relevant to the current study, where we observe that likeability ratings are represented in the BVS during the “distant” condition of the FGAT (where we instructed participants to give a creative association) but not during the “first” condition (where we instructed participants to give the first association that came to their mind). Notably, during the likeability rating task, the instruction was to rate the associations in the context of the FGAT-distant condition (where the goal was to be creative). As a result, the ratings given by the participants may reflect goal values more than subjective values. If so, since the goal to think creatively is similar in the FGAT-distant condition and the likeability rating task, the goal value would remain the same, which might explain why we observed its encoding in the BVS. In contrast, the goal in the FGAT-first condition does not involve creativity and thus differs from that of the likeability rating task. This could explain our null result regarding the correlation between the brain activity during the FGAT-first and the likeability ratings, as these would reflect the goal value for a goal that has yet to be established.

Some studies also linked the BVS to the encoding of yet other dimensions. They observed that novelty processing was related to increased activity within the ventral striatum (B. C. Wittmann et al., 2008), the substantia nigra and the ventral tegmental area (Bunzeck & Düzel, 2006; Schott et al., 2004; B. C. Wittmann et al., 2007, 2008). This heightened activity was viewed as a signal for learning when it was associated with enhanced synaptic plasticity in the hippocampus (Düzel et al., 2010). However, these studies did not assess the relationship between BVS and novelty processing with creative functions.

### The role of the BVS in creativity

Despite the widely acknowledged impact of motivation on creativity, the role of the BVS has been largely overlooked, with only a few studies debating its involvement in creativity, mainly for originality or adequacy processing. Huang et al. (2015) showed that, during creative evaluation, BVS regions, such as the caudate nucleus and the substantia nigra, represented originality. In a later study, however, they showed that other regions from the BVS, namely the ventral striatum and orbitofrontal cortex, actually represented adequacy (Huang et al., 2018). Similarly, Matheson et al. (2023) found that medial prefrontal cortex (mPFC) regions which belong to both the BVS and the DMN supported adequacy processing. Additionally, functional connectivity in subcortical regions of the BVS, such as the putamen (Gao et al., 2021) and grey matter volumes in the striatum and ventral tegmental area (Beversdorf, 2019; Takeuchi et al., 2010) were all related to better divergent thinking abilities.

The involvement of the BVS in creativity is corroborated by investigations at the molecular level. Several studies have highlighted that dopaminergic modulation of frontostriatal networks could support creative cognition: the release of dopamine in the striatum could enhance creativity by enabling cognitive flexibility, and in the prefrontal cortex, dopaminergic modulation could help with the need for persistence in creativity (Boot et al., 2017; Zabelina et al., 2016). Another study showed an inverted U-shape effect of dopamine on creativity, which builds on the notion that creative abilities rely on an optimal trade-off level of dopamine (Chermahini & Hommel, 2010). However, this study’s proxy for subcortical dopaminergic function (the eye blink rate) does not differentiate between the different dopamine pathways and receptors. Similarly, Aberg et al. (2017) showed that individuals with decreased dopamine in the right hemisphere scored better in creativity tasks requiring remote associations (Alternative Uses Task and Remote Associates Test).

An overarching complication when studying motivation and creativity is the anatomical similarity between the BVS and the DMN. In the current study, we assessed the overlap of the DMN (as defined by Yeo and colleagues’ seven networks atlas (2011)) and the BVS (as defined by the “reward” term-based meta-analysis of the Neurosynth platform, https://neurosynth.org/, Yarkoni et al., 2011) and quantified a total of 15 671 voxels, a volume which amounts to 6% of the DMN, or 14% BVS (Table S1). This anatomical similarity can be a common issue in the literature where some creativity studies label “DMN” regions that would be labeled “BVS’’ by neuroeconomists. It will be critical for future research to be aware of this overlap and eliminate this source of confusion, for instance, by using localizers, as we did in the current study.

Regardless, while the literature does not clearly establish how the BVS and its related dopaminergic pathways are involved in creativity, it converges towards confirming their central role (Aberg et al., 2016; Oh et al., 2020). Nevertheless, to the best of our knowledge, the present study is the first one to demonstrate the involvement of the BVS in encoding an idea’s subjective value during idea evaluation and production. Previous findings linking the BVS with originality or adequacy processing may only be due to the correlation of likeability with these two dimensions. In contrast, our innovative design helps break down idea evaluation into the monitoring of ideas’ originality and adequacy and the valuation of ideas, clarifying the specific role of the BVS in this process. We also challenge the established roles of the DMN and ECN by investigating the encoding of originality and adequacy, whose neural correlates remain unclear.

### The roles of the DMN and ECN in creativity: revisiting the classical generative and evaluative distinction

The current literature broadly establishes that the DMN is involved in the generation phase of creativity in a spontaneous manner that helps form new associations. In parallel, the ECN is reported to support idea evaluation and selection in a goal-directed and controlled manner. This one-to-one matching of the dual process of creativity to a dual-network organization is supported by many studies suggesting that the DMN is involved in idea generation (Liu et al., 2015; Lloyd-Cox, Chen, et al., 2022; Matheson et al., 2023; Mayseless et al., 2014, 2015; Wang et al., 2023); that the ECN is involved in idea evaluation (Huang et al., 2018; Liu et al., 2015; Lloyd-Cox, Chen, et al., 2022; Matheson et al., 2023; Mayseless et al., 2014; Wang et al., 2023; Wu et al., 2022); and that creativity relies on these networks’ cooperation (Beaty et al., 2019; R. Beaty et al., 2015, 2016, 2018; R. E. Beaty, Cortes, et al., 2021; Kleinmintz et al., 2019).

However, some findings challenge the exact mapping of the two-fold view of creativity to its dual neural basis. First, several studies, like ours, report that idea evaluation can be partly supported by the DMN (Ellamil et al., 2012; Liu et al., 2015, 2015; Lloyd-Cox, Chen, et al., 2022). Some, even more in alignment with our results, precise that the DMN is related to originality processing (Matheson et al., 2023; Mayseless et al., 2014; Wu et al., 2022). If we consider that achieving originality requires making new associations, it makes sense that the DMN, which has consistently been associated with associative processes during generation, would encode this dimension during evaluation. However, other studies slightly diverge from our own and find that the DMN supports the processing of adequacy (Huang et al., 2015). Despite some diverging results that could be due to tasks and analysis differences - many studies challenge the now common conception that the DMN is only involved in idea generation. Second, other studies, like ours, find that the involvement of the ECN is indeed related to the evaluation of an idea, particularly its adequacy dimension (Matheson et al., 2023; Wu et al., 2022). If we consider that achieving adequacy requires exploring candidate ideas and sorting through them to find the most adequate one, it makes sense for this to be associated with cognitive control processes and, therefore, supported by the ECN. However, contrary to our results, others find that the ECN supports the evaluation of an idea’s originality (Huang et al., 2018; Matheson et al., 2023; Mayseless et al., 2014).

In summary, although the studies cited above lack perfect convergence, they question the neural bases of the dual process model of creativity and suggest that evaluation (as in our study) processes rely on both the DMN and ECN.

### The complementary roles of the BVS, the DMN and the ECN in creativity

Beyond their brain location, more and more studies question the computations at play during creativity, particularly its evaluation phase (Huang et al., 2018; Kleinmintz et al., 2019; Lin & Vartanian, 2018). They highlight that this step is often simplified and overlooks the importance of emotions, motivation or valuation. Based on Huang et al.’s (2018) results, which showed that originality and adequacy processing relied respectively on regions of the ECN and the BVS, Kleinmintz et al. (2019) attempted to dissociate evaluation into “monitoring” and “valuation” subprocesses. They propose that the assessment of originality requires cognitive control akin to monitoring and that, in contrast, the assessment of adequacy is linked to emotional and motivational processes akin to valuation.

Our view partly differs from theirs. First, we dissociate evaluation into monitoring and valuation as well but align monitoring with the evaluation of adequacy and originality (i.e., the monitoring of the dimensions necessary for a creative goal) and valuation with the assignment of a subjective value that combines the monitored adequacy and originality.

Second, we show that the ECN encodes adequacy, not originality and that the DMN encodes originality. Finally and most importantly, we show that the BVS underlies the valuation of ideas, not the processing of adequacy. These diverging results may stem from variations in task design: in the study conducted by Huang et al. (2018), participants reviewed creative solutions to riddles and assessed whether they were adequate or not, while here, participants reviewed creative word associations and assessed how much they liked them, and how adequate and original they were. Additionally, methodological differences might contribute to divergences in results: Huang et al. (2018) used high versus low adequacy and originality contrasts, while we used parametric modulation based on adequacy and originality ratings. Overall, we agree with Kleinmintz et al. (2019) that what is commonly labeled evaluation should be subdivided into monitoring and valuation subprocesses. We propose an alternative model in which originality and adequacy are monitored and integrated during valuation, which then guides the final selection step during idea production.

A remaining important question is whether evaluation’s monitoring and valuation subcomponents are consistently both at play during creativity. In the current study, we show that both valuation and monitoring of originality and adequacy are at play during the production of creative word associations: it would be interesting to generalize these results to other creativity tasks. This would require creativity studies to include likeability rating tasks in their design to assess the participants’ subjective values and check if they correlate with behavioral and neural variables (such as the response times and the BVS activity used in the current study).

Notably, the current study finds that subjective value scales with originality and adequacy ratings. First, this result means that valuation builds on originality and adequacy, the two key dimensions of creativity. Second, it adds to prior research on multi-attribute value-based decision-making. This literature studies choices that depend on the value of several criteria; for example, food choices can depend on taste and health attributes. However, the brain networks implicated in multi-attribute value-based decision-making are still unclear. Indeed, the neural representation of attributes may be contingent upon their nature, and in some cases, attributes may not be represented at all in the brain. For instance, in the context of food choices, regions such as the dorsolateral prefrontal cortex (dlPFC) are associated with the healthy attributes of food, while the ventromedial prefrontal cortex (vmPFC) is linked to the tasty attributes (Hare et al., 2011; Lee & Hare, 2023). Similarly, in the realm of social decision-making, brain regions like the temporoparietal junction (TPJ) or the medial prefrontal cortex (mPFC, part of the DMN) are implicated in processing social attributes, while the ventral striatum plays a role in self-related attributes (Matsuura et al., 2021; Ratcliff & McKoon, 2008; Suzuki et al., 2017; M. K. Wittmann et al., 2016). Yet, the neural representation remains uncertain for risky choices and perceptual attributes like shape or colors, with some evidence for attribute representation in the dlPFC and combined value representation in the vmPFC (Kahnt et al., 2011). Notably, several studies do not report specific activity directly linked to attributes. Instead, some suggest that attributes are represented in the same areas as the combined value (Magrabi et al., 2022). Certain studies, such as the one by Hunt et al. (2014), have focused on comparing attributes in the intraparietal lobule, while others propose the involvement of the dlPFC, a component of the ECN, in attribute representation and comparison (Busemeyer et al., 2019). It is essential to highlight that these studies often report brain activities in specific regions rather than brain networks, making direct comparisons challenging. For example, the TPJ and mPFC, both part of the DMN, substantially overlap with the social network (Mars et al., 2012).

Here, we find that during creative valuation, the attributes of creative ideas (originality and adequacy) are separately represented in networks consistently reported in creativity research (in the DMN and ECN, respectively). These findings illustrate that attributes can be represented in different brain regions and that these regions (here, creativity networks) could be relevant to the specific nature of the attributes under consideration (here, creative dimensions).

Finally, the significant correlation between the neural task-related parameter associated with DMN and ECN activities (α_neural_) and the behavioral parameters related to originality and adequacy assessments (α_behavioral_) suggests a neural basis for individual preferences in creativity. The persistence of this correlation during a resting-state session highlights the enduring functional connectivity of these neural networks. These findings underscore the dynamic interplay between cognitive networks involved in creativity and subjective valuation, offering valuable insights into the neural underpinnings of preferences and decision-making in creativity.

### The link between idea valuation and creative abilities

The present study not only shows that individuals combine the dimensions of originality and adequacy into a subjective value, but it also demonstrates that the way they combine them (as estimated by the α and δ parameters of the CES model) relates to their creative abilities: giving more weight to an idea’s originality over its adequacy (i.e., having a high α) and preferring a trade-off of these two dimensions rather than an extreme of either (i.e., having a low δ) correlated with higher scores in creativity tests.

This finding replicates Lopez-Persem et al. (2023) and suggests, together with other studies, that the evaluation step is essential in the creative process. For instance, previous results showed that an overly stringent evaluation inhibited creativity (Benedek et al., 2016; Mayseless et al., 2014). It was also illustrated by Kleinmintz et al.’s study (2014), in which they found that improviser musicians were more lenient than non-improviser musicians when rating creative ideas and that this helped them achieve better creativity.

In parallel, Diedrich et al. (2015) showed that scores in a creativity task were correlated to the idea’s originality and not to its adequacy. Adequacy did predict creativity but only in very original ideas, as the interaction of originality and adequacy explained additional variance of the creativity scores. Here, we computationally quantify similar effects with the α and δ parameters: creativity is linked to giving greater importance to originality (high α) but still preferring a trade-off of originality and adequacy over extremely original but poorly adequate options (low δ).

### Limitations and perspectives

The present study identifies that likeability, originality and adequacy are respectively encoded in the BVS, DMN and ECN as part of an evaluation process, both during a rating task and a creative production task. However, these interpretations are based on parametric modulation analyses, which strictly indicate brain regions where the BOLD signal increases as a function of a given factor (the parametric modulator) that varies across trials. Instead of concluding that “the likeability of ideas is encoded in the BVS during the rating task”, a more conservative interpretation would be “the greater the likeability rating, the greater the BOLD signal in regions of the BVS”. We can thus not exclude that another variable might mediate these correlations.

In other studies, higher activity in the BVS during evaluation was interpreted as emotional arousal (Huang et al., 2015) or motivation (Liu et al., 2015). Additionally, Wu et al. (2022) proposed that higher activity in the DMN during evaluation may stem from participants generating alternatives while viewing the one in front of them. Our analyses cannot rule out these alternative explanations. For instance, generating original ideas might require more activity in the DMN without necessarily implying the DMN’s direct involvement in the evaluation step of creativity. However, the fact that this pattern also emerged in the rating task strongly supports the latter possibility.

### Conclusion

In the current study, we clarify some cognitive and neural mechanisms involved in idea evaluation, a critical but long-overlooked step of the creative process. We combined creativity and neuroeconomics methods to show that valuation participates in idea evaluation, meaning that individual preferences influence the creative process. These preferences depend on a trade-off of originality and adequacy, which are primary creativity criteria. At the neural level, we add the BVS to the main creativity networks classically reported, the ECN and DMN.

Future research should focus on how these three networks interact during idea evaluation. In parallel, to delve deeper into the decision-making processes of creativity, it would be helpful to investigate the selection step of creativity, i.e., how individuals choose the best candidate idea. Like monetary or food choices, creative choices are likely driven by subjective values, as Lopez-Persem et al. (2023) suggested. This also paves the way for investigating suboptimal or biased choices, which are a central issue considering the growing literature on cognitive biases in creativity research (Benedek et al., 2016; Eling et al., 2015; Howard-Jones, 2002; Mueller et al., 2012; Rietzschel et al., 2010; Zhu et al., 2017).

## 3. Material and Methods

### 4.1. Participants

An official ethics committee approved the study (CPP Ouest II – Angers). We recruited and tested forty participants at the CENIR platform of the Paris Brain Institute (ICM). All participants were French native speakers, right-handed, with correct or corrected vision, and no neurological or psychological disease history. We excluded two participants from analyses, one because we interrupted MRI scanning after they suffered from a claustrophobic episode and one because of a technical issue during MRI scanning. The final sample comprised thirty-eight participants 19 females, age = 26.5 ±0.7 (M±SEM), years of education = 16.7 ±0.4 (M±SEM)). They were paid 110€ for a four-hour testing session that included MRI scanning and cognitive and creativity tests inside and outside the scanner.

### 4.2. Experimental design

After giving informed consent, participants completed an MRI session composed of an anatomical T1 scan, three task-based fMRI scans while performing an FGAT, a rating task and a choice task, and a resting-state fMRI scan. Participants then completed an extra rating task and a battery of creativity tests outside the MRI. We used Matlab (MATLAB. (2020). 9.9.0.1495850 (R2020b). Natick, 624 Massachusetts: The MathWorks Inc.) to program the tasks and the Qualtrics software (Qualtrics, Provo, UT, USA. https://www.qualtrics.com) to implement creativity tests.

#### 4.2.1. Free Generation of Association Task

In the Free Generation of Association Task (Figure 1B), which is reported to capture critical aspects of creativity, participants generate creative word associations (Bendetowicz et al., 2018; Prabhakaran et al., 2014). In our experimental design, the task comprised two successive conditions, each comprising 5 training trials and 62 randomized test trials. Each trial displayed a fixation cross for 1.6 to 3.2 seconds, jittered with a uniform distribution. Then, a cue word was displayed. In the “first” condition, participants had up to 10 seconds to give the first word that came to mind after reading the cue word (for example, participants might answer “father” to the cue word “mother”). In the “distant” condition, they had up to 20 seconds to provide a distant word, so the cue and response words would result in a creative word association (for example, participants might answer “nature” to the cue word “mother”). Participants pressed the response button with their right index finger when they had an answer in mind. Then, they said the response word out loud into a microphone. The experimenter typed the response and displayed it on the participant screen. Participants then had the opportunity to repeat their response in case the experimenter had misheard (by pressing the response button again with their right index finger) or to validate the response (by pressing the validation button with their right ring finger). The exact instructions given to the participants are in the *Supplementary Methods*.

#### 4.2.2. Rating task

##### Selection of the FGAT associations for the rating tasks

We split the rating task in two (a likeability rating task performed inside the scanner and an originality and adequacy rating task performed outside the scanner). Each rating task counted five training trials and a variable number of test trials (M±SEM=145±2), depending on the validity of their FGAT answers (see *Supplementary Methods* for more details). For both rating tasks, the trials were composed of the following: 66% of the trials were evenly made up of the participant’s valid FGAT-first and FGAT-distant associations; 27% of the trials were a sample of frequent and rare responses taken from another study using the same FGAT cue words; and finally 7% of the trials were responses that were totally unrelated to the cue words they were associated with (for example: ‘cow’ and ‘inverse’). We added these associations to cover a wide range of adequacy and originality ratings, ensuring a robust estimation of likeability with sufficient statistical power.

##### Likeability rating task

The first rating task, i.e. the likeability rating task (Figure 1C), was performed in the MRI scanner. Participants had to indicate how much they liked the association or how satisfying it was in the context of the FGAT-distant condition. In this task, each trial started with a fixation cross displayed for 1.6 to 3.2 seconds (display times were determined by a random jitter with a uniform distribution). Then, a cue word (for example, “mother”) was displayed alone for 0 to 0.5 seconds (display times were determined by a random jitter with a uniform distribution). Then, a response appeared under the cue word. This association (for example, “mother-nature”) remained alone for 1.6 to 3.2 seconds (display times were determined by a random jitter with a uniform distribution). Then, a rating scale appeared below the association. The rating scale’s low to high values were represented from left to right, with the words “not at all” and “very much” located at the extremities but without any numerical values. A segment indicated the middle of the scale, which was divided into 101 hidden steps, later converted into ratings ranging between 0 and 100. We displayed a heart icon (for likeability) under the rating scale. To provide a rating, participants had to move a cursor by pressing the first and second MRI response buttons with the right index and right middle finger and then validate the rating by pressing the third button with their right ring finger. Once the response was validated, the subsequent trial began. The exact instructions given to the participants are in the *Supplementary Methods*.

##### Originality and adequacy rating task

The second rating task, i.e. the originality and adequacy rating task (Figure 1D), was completed after the choice task and outside the MRI. Again, each trial displayed a two-word association (for example, “mother-nature”) and asked participants to subsequently rate how original and how adequate it was. The order of originality and adequacy ratings was counterbalanced across trials. Each trial started with the screen displaying a cue word. After 0.3 seconds, a response word appeared under the cue, and after 0.4 seconds, a scale appeared under the two-word association. The scale was similar to the likeability rating scale except for the icon displayed underneath: a star for originality and a target and arrow for adequacy. Participants rated the association by moving the slider across the scale using the left and right arrows of the keyboard and validated by pressing the spacebar. Then, a new scale appeared underneath the same association, indicating the second dimension (adequacy or originality). Participants provided their ratings similarly and then started the subsequent trial.

We told participants to use the whole scale throughout the task. No time limit was applied, but the instruction was to respond spontaneously. The exact instructions given to the participants are in the *Supplementary Methods*.

#### 4.2.3. Choice task

The choice task was performed in the MRI and comprised five training trials and an average of 187 randomized test trials (the exact number varied between participants depending on the variability of their FGAT answers and ratings). Each trial displayed two possible responses to an FGAT cue, with the cue word on the top of the screen and the two response options words below, on the same horizontal line (for example, “mother” was displayed on top of “nature” and “father”). Participants had to choose which of the two responses would result in the best association with the top cue in the FGAT-distant condition. No time limit was applied, but the instruction was to respond spontaneously. Note that the results of this task go beyond the scope of this paper and will instead be the focus of a future publication. The exact instructions given to the participants are in the *Supplementary Methods*.

#### 4.2.4. Battery of creativity tests

The battery comprised a daily-life creativity questionnaire, self-reports, and established creativity tasks belonging to three main frameworks of creativity - the divergent thinking approach, the associative theory, and the insight problem-solving approach.

<u>Inventory of Creative Activities and Achievements:</u> the inventory of creative activities and achievements was first designed by Diedrich et al. (2018) as an ecological measure of creativity. It focuses on eight domains of creativity (literature, music, crafts, cooking, sports, visual arts, performing arts, and science and engineering). For each domain, participants reported their level and their frequency of engagement over the past ten years. They also specified their level of creative accomplishment in each of these domains.

<u>Scoring:</u> we computed the ICAA score as the mean of the creative activities score (C-act) and the creative achievements score (C-ach). First, for the creative activities score (C-act), we summed the frequency at which the participant had engaged in each of the eight creativity domains using an ordinal scale ranging from 0 (never) to 4 (more than ten times). Then, for the creative achievements score (C-ach), we summed the participants’ levels in each of the eight creativity domains. These ranged from 0 (“never engaged in this domain”) to 10 (“I have already sold some of my work in this domain”).

<u>Drawing task:</u> the drawing task (Barbot, 2018) comprised a training trial and 12 test trials. We instructed participants to include incomplete shapes in creative drawings. The 12 incomplete shapes were relatively similar: they were all composed of 4 lines, either straight, curved or right-angled, and exhibited horizontal symmetry (cf. (Barbot, 2018) for an example). Participants completed the 12 drawings at their own pace. Then, they reviewed each drawing and gave each one a verbal title.

<u>Scoring:</u> We followed the consensual assessment technique that is advised for scoring such a creativity task (Ceh et al., 2022). We designated four judges from the lab who were familiar with creativity judgments but had not seen the data before the rating. For each of the twelve shapes, the judges first saw the incomplete shape, then each participant’s drawing using that shape, in a blind and randomized order. Then, the judges gave each drawing a rating between 0 (not creative at all) and 4 (extremely creative). We tested the inter-judge independent rating consistency using an intraclass correlation analysis, which gives the mean correlation between the judges’ ratings (r=0.8239 ± 0.02; M±SEM).

<u>Combination of associates task</u>: Bendetowicz et al. (Bendetowicz et al., 2018) first developed the combination of associates task by adapting the remote associative task from Mednick (1962). After five training trials, participants performed 40 randomized test trials, each displaying three cue words with no apparent association: they had to find a fourth word that linked all of them (for example, the link between “map”, “dig”, and “discover” is “treasure”).

Scoring: The CAT score used for the analyses was the participants’ total number of accurate answers.

<u>Associative fluency task</u>: in this task, adapted from Benedek et al. (2012), participants had two minutes to type as many words related to a cue word as possible. We chose this associative design over simple category or letter fluency tasks to measure creative abilities on top of fluency. We selected six cue words (”garden”, “wine”, “rock”, “opinion”, “call”, and “finger”) among the FGAT cues for their diversity in steepness. The steepness is the ratio of the frequency of the most frequent answer over the frequency of the second most frequent answer: a steep cue word (e.g., “cat”) has a very common first associate (“dog”); a flat cue word has several equally-common first associates. We measured typing speed and accounted for it in the fluency analysis.

Scoring: The score used for the analyses was the total number of answers given to a cue word, averaged across all cue words.

Participants also performed an alternative uses task (AUT), a self-report of their creativity level (i.e. “How creative are you on a scale of 1 to 100?”) and a questionnaire regarding their preferences in creativity (e.g., “Would you prefer an idea to be useful or novel?”) that were not analyzed or used in this paper but will be part of future papers.

### 4.3. Behavioral analyses

We performed all the analyses using Matlab (MATLAB. (2022). 9.12.0.2009381 (R2022a). Natick, Massachusetts: The MathWorks Inc.).

#### 4.3.1. Differences in originality, adequacy and frequency between FGAT-first and FGAT-distant responses

For each individual, we computed the mean of originality and adequacy ratings separately for items of the FGAT-first and FGAT-distant conditions. Then, we compared the difference in adequacy between the two tasks to their difference in originality using a paired two-tailed t-test at the group level. We computed the frequency of association of all FGAT responses using the French database Dictaverf (http://dictaverf.nsu.ru/) (Debrenne, 2011). This database is built on spontaneous associations provided by at least 400 individuals in response to 1081 words (each person saw 100 random words). Frequencies were log-transformed to take into account their skewed distribution toward 0. We compared the frequency of the FGAT-first responses and the FGAT-distant responses at the group level using a paired two-tailed t-test.

#### 4.3.2. Relationship between FGAT response times and likeability ratings

We defined participants’ response times in the FGAT as the time between the cue display and the participants’ first button press, which indicated they had an answer in mind. Then, at the individual level, we performed a linear regression of response times against likeability ratings, standardized between participants. We did two distinct regressions for the FGAT-first and FGAT-distant conditions. At the group level, we performed a one-sample, two-tailed t-test to test the significativity of the effect of likeability on response times. We also tested the difference in the effect of likeability on response times (regression coefficients) between the two FGAT conditions using a paired two-tailed t-test on the regression coefficients. Note that all variables were standardized at the subject level, and non-standardized data are depicted in figures.

We performed a control analysis to compare the effects of likeability and confidence on response times. We did a GLM on response times at the individual level, where we used likeability ratings and squared likeability ratings (a proxy for confidence) as regressors. We standardized response times and likeability ratings between participants. We compared the effects of likeability ratings and squared likeability ratings at the group level using a paired two-tailed t-test.

#### 4.3.3. Relationship between likeability, originality and adequacy

We fitted the participants’ data at the individual level with the Constant Elasticity of Substitution utility function (Andreoni & Miller, 2002; Lopez-Persem et al., 2017), which provided the best fit in our previous study using the same experimental design (Lopez-Persem et al., 2023):

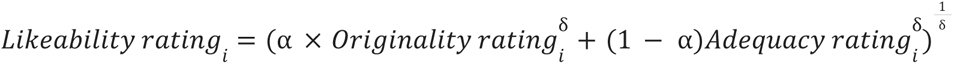

“i” stands for one association, α stands for the weight given to originality ratings, and δ stands for the preference for extremes in originality or adequacy as opposed to a trade-off of those dimensions.

We used the Matlab VBA toolbox (https://mbb-team.github.io/VBA-toolbox/), which implements Variational Bayesian analysis under the Laplace approximation (Daunizeau et al., 2009; Stephan et al., 2009) to fit the model to individual data and estimate individual parameters. We tested for significance the estimated parameters of the CES model at the group level using paired two-tailed t-tests against their prior values (0.5 for α and 1 for δ).

#### 4.3.4. Post-hoc analyses on the alpha parameter and the relationship between likeability and frequency

Next, we performed post-hoc analyses on the α parameter of the CES, which captures the weight of originality relative to adequacy in the likeability ratings. First, we computed the coefficient of variation: α’s standard deviation divided by its mean value.

In parallel, we performed a linear regression of likeability against frequency at the individual level on standardized values. We tested its significativity at the group level using a one-sample, two-tailed t-test. Then, we tested the group-level correlation between the α parameter and the slope of the likeability-frequency linear regression.

#### 4.3.5. Creative abilities analyses and canonical correlation

We pooled the scores for the associative fluency task, the Combination of Associates Task (CAT), the drawing task and the Inventory of Creative Activities and Achievements (ICAA) (see *Material and Methods 4.2.4* for more information) into a “creativity scores” set and the α and δ parameters of the evaluation model into an “evaluation parameters” set. Then, we performed a canonical correlation analysis of these two sets. Canonical correlation analysis identifies the shared variance between two datasets, summarizing it as canonical variables. They are organized based on the strength of correlations between the two datasets. Here, we examined the correlation between the canonical variables in each dataset and presented their loading coefficients, i.e. the contributions of each creativity score and each evaluation parameter to their respective canonical variable. Correlation coefficients were tested for significance using Bartlett’s modified chi-squared tests of the Matlab “canoncorr” function, and loading coefficients were tested for significance using the two-tailed t-test of the Maltab “corr” function.

### 4.4. MRI data acquisition and preprocessing

#### 4.4.1. Scanning parameters

We acquired neuroimaging data on a 3T MRI scanner (Siemens 3T Magnetom Prisma Fit) with a 64-channel head coil.

Four functional runs were acquired over the four tasks (FGAT-first, FGAT-distant, likeability rating task, choice task) using multi-echo echo-planar imaging (EPI) sequences. The number of whole-brain volumes per run varied between tasks but also between participants since all tasks were self-paced (mean, [min; max]: FGAT-first= 379, [317;454], FGAT-distant: 612, [359;856]; likeability rating task: 800, [650;1012] and choice task: 662, [525;821]). We did not record any dummy scans and, therefore, did not discard any volume. The functional runs used the following parameters: repetition time (TR) = 1660 ms; echo times (TE) for echo 1 = 14.2 ms, echo 2 = 35.39 ms, and echo 3 = 56.58 ms; flip angle = 74°; 60 slices, slice thickness = 2.50 mm; isotropic voxel size of 2.5 mm; Ipat acceleration factor = 2; multiband = 3; and interleaved slice ordering.

After the EPI acquisitions, we acquired a T1-weighted structural image with the following parameters: TR = 2300 ms, TE = 2.76 ms, flip angle = 9°, 192 sagittal slices with a 1-mm thickness, isotropic voxel size of 1 mm, Ipat acceleration factor = 2.

Finally, we added a resting-state fMRI session of 10 min (360 volumes) with the same acquisition parameters as the task runs.

#### 4.4.2. Preprocessing

We performed the preprocessing of the on-task fMRI data separately for each run and the resting-state data using the afni_proc.py pipeline from the Analysis of Functional Neuroimages software (AFNI; https://afni.nimh.nih.gov). The different preprocessing steps of the data included slice timing correction and realignment to the first volume (computed on the first echo). We then combined the preprocessed data using the TE-dependent analysis of multi-echo fMRI data (TEDANA; https://tedana.readthedocs.io/), version 0.0.9a1 (DuPre et al., 2021; Kundu et al., 2013). We co-registered the resulting data on the T1-weighted structural image using the Statistical Parametric Mapping (SPM) 12 package running in MATLAB (MATLAB R2017b, The MathWorks Inc., USA). We normalized the data to the Montreal Neurological Institute template brain using the transformation matrix computed from the normalization of the T1-weighted structural image with the default settings of the computational anatomy toolbox (CAT 12; http://dbm.neuro.uni-jena.de/cat/) (Gaser et al., 2022) implemented in SPM 12.

### 4.5. fMRI analyses

#### 4.5.1. Parametric modulation analyses

We entered the resulting normalized data from the task-based fMRI in general linear models in SPM. In this analysis, we entered 24 motion parameters and 42 physiological noise parameters as confounds regressed from the BOLD signal. In this preprocessing analysis, we used the Matlab PhysIO Toolbox (Kasper et al., 2017) (version 5.1.2, open-source code available as part of the TAPAS software collection: https://www.translationalneuromodeling.org/tapas (Frässle et al., 2021)) to generate nuisance regressors for the GLM. The regressors were composed of (i) physiological (cardiac pulse and respiration) recordings used to generate RETROICOR (Glover et al., 2000) regressors, (ii) White Matter (WM) and CerebroSpinal Fluid (CSF) masks from the anatomical segmentation were used to extract components from compartments of non-interest using Principal Component Analysis (PCA) and (iii) motion parameters, composed of standard motion parameters, first temporal derivatives, standard motion parameters squared, and first temporal derivatives squared. Outlier volumes were eliminated when the Framewise Displacement (FD) exceeded 0.5mm.

Then, we used GLMs to explain pre-processed time-series at the individual level.

<u>Modelling the neural encoding of likeability during idea evaluation (functional localizer):</u> We applied the first, second and third models (GLM1, GLM2 and GLM3) to the likeability rating task. They all included a boxcar function capturing the signal between the cue display and the first button press to move the rating scale slider (one event per trial). GLM1’s parametric modulator was the likeability rating of the response, while GLM2 and GLM3 were parametrically modulated by the originality and adequacy ratings, respectively. Note that GLM2 and GLM3 parametric modulators (originality and adequacy ratings, respectively) were not orthogonalized for the other dimension (meaning adequacy and originality, respectively), but control analyses with orthogonalization led to similar results.

<u>Modelling the neural encoding of likeability during idea production:</u> The fourth, fifth and sixth models (GLM4, GLM5 and GLM6) were similar to the first three models, except that we applied them to the FGAT-distant. They all included a boxcar function capturing the signal between the cue display and the first button press, signaling that the participant had found a response to the cue (one event per trial). GLM4’s parametric modular was the likeability rating of the response, while GLM5’s and GLM6’s were the originality and adequacy ratings, respectively. Note that not all 62 FGAT-distant trials were included in this analysis: depending on the participant, the number of trials ranged between 34 and 59, as some associations had not been rated after controlling for FGAT-first and FGAT-distant similarity (see *Supplementary Methods* for more details).

We convolved the regressors for all models with a canonical hemodynamic response function. The resulting masks were identified in statistical parametric maps (SPMs) at a threshold of p < 0.05, controlling for family-wise errors (FWE) at the cluster level. For GLMs 1 to 3, we found significant clusters at the whole brain level. For GLMs 4 and 5, we found significant clusters using small volume correction (SVC) with the overlap of the localizer and the networks of interest. For GLM6, SVC were spheres centered on the peak activation sites from the results of GLM3.

#### 4.5.2. Functional localizers’ similarity with networks of interest

To compare our resulting maps to our networks of interest, namely the BVS, the DMN and the ECN, we quantified the number of overlapping voxels between the SPMs of GLMs 1 to 3 and each of Yeo et al. (2011) 7 intrinsic functional networks in the MNI152 referential. Since Yeo’s atlas does not include the BVS, we used the term-based meta-analysis of the Neurosynth platform (https://neurosynth.org/, Yarkoni et al., 2011): the “atlas BVS network” is therefore the result of an automated meta-analysis of all studies in the Neurosynth database whose abstracts include the term “reward” at least once.

Before quantifying the overlaps, we verified that the atlas masks aligned with our results using the SPM checkReg function and applied it to our data with the ImCalc tool in SPM12.

#### 4.5.3. Fitting the CES to the FGAT-distant and resting-state timeseries

To investigate the interaction between the BVS, ECN and DMN during the FGAT-distant, we employed a procedure previously used in the lab (Ovando-Tellez et al., 2022). To covary out the task-related signal from the FGAT distant run, we entered the preprocessed fMRI data in a GLM in SPM. We regressed out the onsets of each task-related event (cross onset, cue onset, first press event, validation event) from the BOLD signal. We then standardized and detrended the residuals of the GLM and concatenated them to obtain timeseries for each voxel. Timeseries for all voxels belonging to each network were averaged across voxels. Timeseries from the DMN and ECN were orthogonalized to remove shared variance between them. Using the VBA toolbox, we then fitted the following equation: BVS_FGAT-distant_ = (*α*DMNF_GAT-distant_^*δ*^ + (1-*α*)ECN_GAT-distant_^*δ*^)^1/*δ*^

The estimated parameters from this equation will be referred to as the FGAT-distant α_𝑛𝑒𝑢𝑟𝑎𝑙_, capturing the relative contribution of DMN and ECN activities to the BVS activity during the FGAT-distant, and the FGAT-distant the δ_𝑛𝑒𝑢𝑟𝑎𝑙_, capturing the curvature of contribution of the DMN and ECN activities to BVS activity.

We preprocessed the resting-state BOLD signal in the same way as the task BOLD activity and extracted the timeseries from each network to apply the same analysis and fit the following equation: BVS_RS_ = (*α*DMN_RS_^*δ*^ + (1-*α*)ECN_RS_^*δ*^)^1/*δ*^

The estimated parameters from this equation will be referred to as the resting-state α_𝑛𝑒𝑢𝑟𝑎𝑙_ and the resting-state δ_𝑛𝑒𝑢𝑟𝑎𝑙_.

We then used Pearson correlations to compare each neural parameter, estimated with the timeseries, to its respective behavioral parameter, estimated with the ratings.

## Funding

EV was funded by the “Agence Nationale de la Recherche” grant (number ANR-19-CE37-0001-01) and the “Investissements d’avenir” program (number ANR-10-IAIHU-06). This project received funding from the European Union’s Horizon 2020 Research and Innovation program under the Marie Sklodowska-Curie grant agreement number 101026191. ALP was by the “Fondation des Treilles”. SMR was supported by the “Frontières de l’Innovation en Recherche et Éducation” PhD program, affiliated with Paris Cité University.

## Contributions and acknowledgments

EV, ALP and SMR designed the study. SMR collected the data and performed all analyses. BB performed the pre-processing of the fMRI. ALP provided template scripts for analyses. SMR, ALP and EV wrote the article. All authors reviewed and edited the article.

We thankfully acknowledge the CENIR platform and especially Céline Homo, Stéphanie Anastacio, and Christine Soeung, as well as Declan Grindrod and Alexandre Bacq, for their help during data collection. We thank Baptiste Barbot for letting us use his drawing task and Marcela Ovando-Tellez, Marie Scuccimarra and Ines Maye for rating the participants’ drawings. Finally, we sincerely thank the forty volunteers who participated in this study.

## Competing interests

The authors report no competing interests.

## Notes

https://github.com/smorenorodriguez/CreHack_NeuroValuation

### Competing Interest Statement

The authors have declared no competing interest.

### Summary of Updates

- change of title - changes in the abstract - minor corrections in the results

