## Supplemental for "The human reward system encodes the subjective value of ideas during creative thinking"

**Supplementary**

1. *Supplementary results*

Below are detailed supplementary control analyses mentioned in the main text.

*Individuals provide preferred responses faster: control analyses.*

Importantly, we took into consideration the quadratic relationship that exists between likeability ratings and confidence, as it has been shown that people are more confident in high and low likeability ratings and more uncertain for intermediate ratings (H. Barron et al., 2015; Lebreton et al., 2015). We sought to confirm that the effect of likeability on response times was a marker of valuation and not confidence. To do so, we introduced the squared standardized likeability ratings as an additional regressor alongside the likeability ratings in the general linear model (GLM) of response times. We found that likeability had a stronger impact on response times compared to the squared likeability ratings ($\beta_{likeability}$=-0.17±0.03, t(37)=-6.66, p=8.10$^{-8}$; $\beta_{likeability\textasciicircum 2}$=0.01±0.02, t(37)=0.38, p=0.71; paired two-tailed t-test: t(37)=-6.87, p=4.10$^{-8}$), confirming that the effect of likeability on response times was a marker of valuation and not confidence.

*Locating the neural encoding of likeability during idea production: control analyses of response times neural correlates*

As response times negatively correlated with likeability ratings in FGAT-distant, we conducted a control analysis using a whole-brain parametric modulation approach, with response time as a regressor. During the rating task, the response times were encoded in several regions, namely in the insula ([-44, 0, -4], t(37)=10.81, pFWE<10$^{-16}$) and parietal operculum ([52, -35, 26], t(37)=10.52, pFWE<10$^{-16}$). During FGAT-distant, response times were mainly encoded in the lateral frontal pole ([32, 50, 23], t(37)=7.62, pFWE=2.10$^{-10}$) and the supramarginal gyrus ([62, -48, 28], t(37)=7.28, pFWE=1.10$^{-11}$). We found no significant correlation between response times and the BOLD signal in regions of the BVS during the likeability rating task or FGAT-distant (see Table S2 for all significant clusters).

2. *Supplementary methods*

Below are supplementary method details mentioned in the main text.

*FGAT task instructions*

In the FGAT, we divided instructions into successive slides, illustrated with screenshots of what the task would look like, and participants read them at their own pace. The instructions were as follows (translated from French):

*FGAT-first:* “In each trial, you will see a word appear on the screen. You will have to state the first word that comes to your mind without thinking and as fast as possible. As soon as you have a word in mind, you will need to press your index finger to give your response to the experimenter. The word "Response?" will then appear on the screen: you can say the word out loud to the experimenter. The experimenter will write down the word heard. You must verify that it is indeed the word you said. If yes, press with your ring finger to confirm your answer. If not, press with your index finger to repeat your response. Please note that this is only to confirm that the word has been heard correctly. You do not have the right to change your mind. Once your answer is confirmed, a new word will appear on the screen; proceed in the same way. Be aware! You will have 10 seconds to think of your response. If you take too long to find an answer, we will move on to the next trial. Before pressing with your index finger, wait until you have an answer in mind: you must say it directly to the experimenter without taking additional time to think. Conjugated verbs, proper nouns, and groups of words are not accepted.”

*FGAT-distant:* “Similar to the previous task, with each new trial, you will see a word appear on the screen. This time, you must respond with a word that is not typically associated with the displayed word but is still somehow related. Think creatively: the association between the displayed word and your response should be original, unusual, surprising, while still being understandable to someone else. As soon as you have a word in mind, you will need to press your index finger to give your response to the experimenter. The word "Response?" will then appear on the screen: you can say the word out loud to the experimenter. The experimenter will write down the word heard. You must verify that it is indeed the word you said. If yes, press with your ring finger to confirm your answer. If not, press with your index finger to repeat your response. Please note that this is only to confirm that the word has been heard correctly. You do not have the right to change your mind. Once your answer is confirmed, a new word will appear on the screen; proceed in the same way. Be aware! You will have 20 seconds to think of your response. If you take too long to find an answer, we will move on to the next trial. Before pressing with your index finger, wait until you have an answer in mind: you must say it directly to the experimenter without taking additional time to think. Conjugated verbs, proper nouns, and groups of words are not accepted.”

#### *Rating task instructions*

For the likeability rating task, we divided instructions into successive slides, illustrated with screenshots of what the task would look like, and participants read them at their own pace. The instructions were as follows (translated from French):

“As a reminder, at the beginning of this experiment, you performed two tasks. In response to a word (for example, "jewelry"), you were required to give two types of responses: the first response that comes to your mind (for example, "jewelry-necklace") and an unusual but understandable response (for example, "jewelry-money"). In this task, with each new trial, you will see a pair of words on the screen. The first word is part of the words you have seen in the previous tasks. The second word is a word possibly associated with the first word, which can be a word you proposed in the previous tasks or another suggestion. The task is to indicate how much you like the association between these two words in the context of the previous task (where you had to provide an unusual response). For example, how much would you have liked to respond with the word "necklace" to the word "jewelry" in the previous task? Below the association, an evaluation scale will appear, with a heart. Using this scale, you will indicate how much you like this association. Move the cursor using your index and middle fingers. Confirm your response by pressing with your ring finger. Be aware! You must move the cursor, even if you are perfectly neutral (in this case, move and return the cursor to the middle). Use the entire scale: do not always respond the same way! Reminder: Indicate how much you would have liked to propose the second word of the association in response to the previous task (unusual response). If it's a response you have given before, indicate how satisfied you are with that response.”

For the originality and adequacy rating task, we divided instructions into successive slides, illustrated with screenshots of what the task would look like, and participants read them at their own pace. The instructions were as follows (translated from French):

“As a reminder, at the beginning of this experiment, you performed two tasks. In response to a word (for example, "jewelry"), you were required to give two types of responses: the first response that comes to your mind (for example, "jewelry-necklace") and an unusual but understandable response (for example, "jewelry-money"). Similar to the previous task, with each new trial, you will see a pair of words on the screen. You will evaluate this word association based on two specific criteria. Firstly, does this association seem appropriate to you, meaning understandable, relevant, fitting? To respond, a relevance evaluation scale will appear. Move the cursor using the "left arrow" and "right arrow" keys. Confirm your response by pressing "space." If the association seems entirely relevant to you, move the cursor to the right. If you do not see a connection between the words, move the cursor to the left. Any evaluation between these two extremes is possible. Once your response is confirmed, you will evaluate a second criterion, that of the originality of this word association: how unusual and surprising you find this association. For this, a second evaluation scale will appear. This scale will allow you to indicate how original you think the association is. Move the cursor using the "left arrow" and "right arrow" keys. Confirm your response by pressing "space." Be aware! You must move the cursor, even if you are perfectly neutral (in this case, move and return the cursor to the middle). Try to use the entire scale: do not always respond the same way."

*Rating task trials: selection of the FGAT associations for the rating tasks*

The number of trials in the rating task varied between participants as they saw only associations that followed these criteria:

- the first three letters of the FGAT-first and FGAT-distant responses to the cue word must be different
- one of the responses must not contain the other (e.g. 'count' and 'accountant' have the word 'count' in common and would not respect this criterion)
- the FGAT-first and FGAT-distant responses to one cue word must be comparable in length (no more than five letters of difference)
- the FGAT-distant response must not be too long (no more than nine letters)

Then, we randomly selected one-fifth of the accepted cue words and paired them with five other responses that the participant had not generated: four of those from a different study's dataset, including (i) a highly frequent FGAT-first response, (ii) a highly unfrequent FGAT-first response, (iii) a highly frequent FGAT-distant response and (iv) a highly unfrequent FGAT-distant response. The fifth response was an unrelated word (for example, the cue word 'cow' was associated with the unrelated response 'inverse'). Overall, this added up to a maximum of 186 rating trials (less if the 124 FGAT-first and FGAT-distant associations did not all follow the four criteria mentioned above), selected for their diversity and randomized before the start of the task.

*Choice task instructions*

For the Choice task, we divided instructions into successive slides, illustrated with screenshots of what the task would look like, and participants read them at their own pace. The instructions were as follows (translated from French):
"As a reminder, at the beginning of this experiment, you performed two tasks. In response to a word (for example, "jewelry"), you were required to give two types of responses: the first response that comes to your mind (for example, "jewelry-necklace") and an unusual but understandable response (for example, "jewelry-money"). Similar to the previous task, with each new trial, you will see a word at the top of the screen (for example, "jewelry"), but this time it will be associated with two other proposals (for example, "necklace" and "money"). Choose the association you prefer: "jewelry" with "necklace" or "jewelry" with "money." In other words, would you have preferred to respond with "necklace" or "money" to the word "jewelry" in the second task (unusual response)? To select a

response, use your index finger (left response) or your middle finger (right response). You do not need to validate your choice; it will be framed and validated as soon as you press. Another trial will then start."

*3. Supplementary figures and tables*

Supplementary figures and tables mentioned in the main text are depicted below.

*Individuals prefer responses that are both adequate and original: post-hoc analyses on the alpha parameter.*

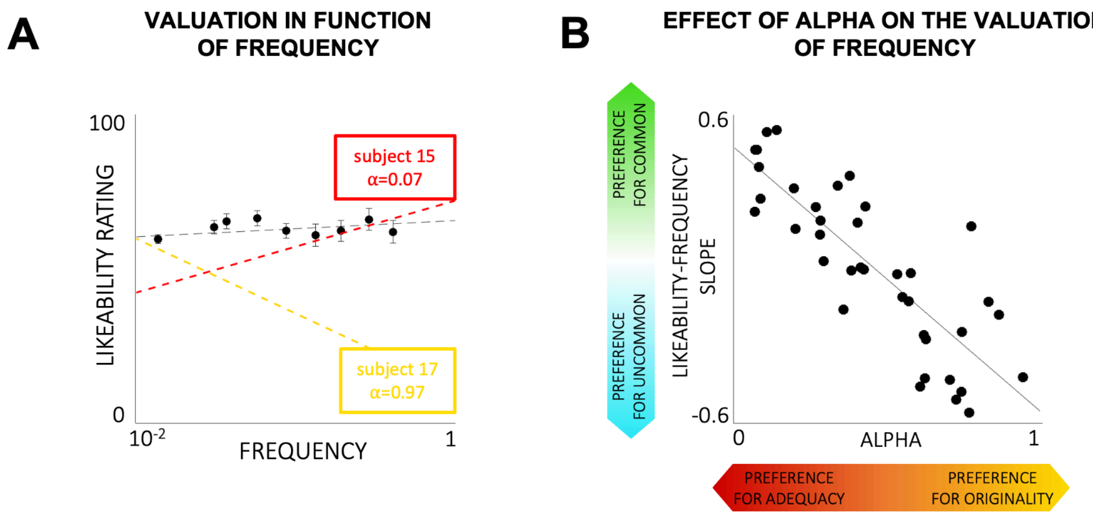


**Figure S1: Link between valuation of frequency and preference for originality. (A)** Correlation between likeability ratings and frequency of associations. Black circles are bins of averaged participant data. Error bars are intersubject standard error of the mean (SEM). The dashed black line indicates a non-significant linear regression fit at the group level ($p>0.05$). To illustrate intersubject variability, the red and yellow dashed lines are linear regressions for two subjects showing respectively a strong preference for adequacy ($\alpha<0.05$) and originality ($\alpha>0.05$). **(B)** Correlation between the likeability-frequency slope of panel A and the preference for originality ($\alpha$). Each circle represents one participant. The solid line indicates a significant correlation at the group level ($p<0.05$).

*Comparing the functional localizer of originality and adequacy evaluation to atlas functional networks : all common voxel counts*

| | likeability ratings | originality ratings | adequacy ratings | BVS | ECN | DMN | visual | somatomotor | dorsal attention | salience | limbic |
|---|---|---|---|---|---|---|---|---|---|---|---|
| likeability ratings | 120078 | | | | | | | | | | |
| originality ratings | 37419 | 236590 | | | | | | | | | |
| adequacy ratings | 26709 | 2035 | 150112 | | | | | | | | |
| BVS | **31688** | 32843 | 5871 | 112370 | | | | | | | |
| ECN (ECN excluding BVS) | 4092 (4025) | 7728 (7704) | **59921** (58713) | 2957 (0) | 172662 (169705) | | | | | | |
| DMN (DMN excluding BVS) | 27718 (19630) | **82262** (77092) | 24227 (22363) | 15671 (0) | 0 | 248106 (232435) | | | | | |
| visual | 17615 | 23842 | 3769 | 18 | 0 | 0 | 150346 | | | | |
| somatomotor | 767 | 188 | 2556 | 40 | 0 | 0 | 0 | 138244 | | | |
| dorsal attention | 2282 | 3346 | 14748 | 0 | 0 | 0 | 0 | 0 | 122876 | | |
| salience | 2322 | 11357 | 5747 | 4955 | 0 | 0 | 0 | 0 | 0 | 131520 | |
| limbic | 10168 | 19756 | 4569 | 12571 | 0 | 0 | 0 | 0 | 0 | 0 | 90979 |

**Table S1: Common voxels in functional networks of interest.** Number of common voxels between the networks of Yeo et al.'s atlas (2011) and the statistical maps of the parametric modulation of the likeability rating task by likeability, originality and adequacy ratings. Numbers in parentheses correspond to the voxel count of the overlap of ECN and DMN networks with BVS regions, which were removed from the DMN and ECN ROIs in the following analyses. Only the results in bold are reported in the main text.

*Locating the neural encoding of likeability during idea production: control analyses of response times neural correlates*

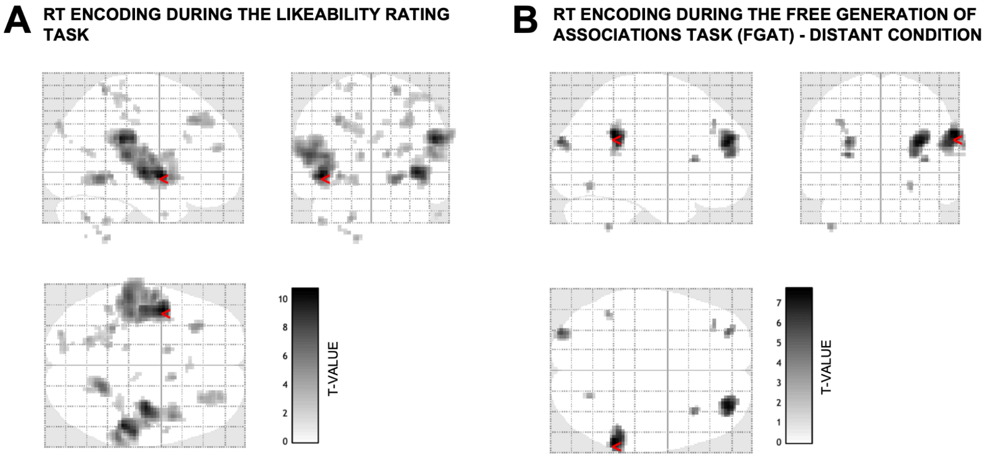


**Figure S2: Neuroimaging results of response times correlates during the likeability rating task and the FGAT-distant. (A)** Neural correlates of RT during the likeability rating task. **(B)** Neural correlates of RT during the FGAT-distant. Shades of grey indicate the T-statistic of one-sample t-tests, FWE $p<0.05$. Red arrows indicate the global maximum. See Table S2 for all significant clusters. Neural correlates do not overlap with the BVS.

| task<br>**likeability rating task** | | | parametric modulator<br>**response time during the likeability rating task** | | | | | |
|---|---|---|---|---|---|---|---|---|
| cluster label | cluster size | peak side | x | y | z | Z-score | T-value | cluster p-value |
| **insula/superior temporal gyrus** | **1248** | **L** | **-44** | **0** | **-4** | **7.18** | **10.71** | **<10-16** |
| insula/superior temporal gyrus | 417 | R | 36 | -15 | -2 | 6.94 | 10.06 | <10-16 |
| **supramarginal gyrus/superior temporal gyrus** | **588** | **R** | **52** | **-35** | **26** | **6.92** | **10.00** | **<10-16** |
| lingual gyrus | 111 | R | 19 | -52 | -7 | 6.22 | 8.35 | 1.10-8 |
| middle frontal gyrus | 45 | L | -31 | 30 | 40 | 5.78 | 7.44 | 1.10-5 |
| lingual gyrus | 83 | L | -28 | -42 | -4 | 5.74 | 7.35 | 1.10-7 |
| superior parietal lobule | 13 | R | 19 | -50 | 66 | 5.70 | 7.28 | 0.001 |
| cerebellum | 17 | L | -18 | -65 | -40 | 5.57 | 7.04 | 6.10-4 |
| caudate nucleus | 21 | R | 12 | 8 | 13 | 5.54 | 6.97 | 3.10-4 |
| middle frontal gyrus | 87 | R | 29 | 32 | 40 | 5.51 | 6.92 | 1.10-7 |
| superior frontal gyrus | 33 | R | 29 | 48 | 18 | 5.44 | 6.79 | 5.10-5 |
| cerebellum | 14 | L | -14 | -48 | -57 | 5.41 | 6.74 | 0.001 |
| caudate nucleus | 20 | L | -11 | 2 | 13 | 5.40 | 6.73 | 3.10-4 |
| occipital cortex | 47 | L | -24 | -88 | 36 | 5.40 | 6.73 | 9.10-6 |
| cerebellum | 19 | L | -46 | -48 | -42 | 5.39 | 6.71 | 5.10-4 |
| postcentral gyrus (S1) | 56 | R | 46 | -32 | 60 | 5.22 | 6.40 | 3.10-6 |
| calcarine sulcus | 16 | R | 22 | -65 | 16 | 5.20 | 6.37 | 0.001 |
| inferior temporal gyrus | 6 | R | 56 | 2 | -37 | 5.16 | 6.30 | 0.006 |
| cerebellum | 9 | L | -21 | -55 | -52 | 5.14 | 6.27 | 0.003 |
| midcingulate cortex | 8 | L | -14 | -32 | 46 | 5.13 | 6.25 | 0.004 |
| superior parietal lobule/postcentral gyrus (S1) | 14 | L | -18 | -50 | 70 | 5.11 | 6.21 | 0.001 |
| anterior cingulate cortex | 16 | R | 4 | 20 | 28 | 5.07 | 6.15 | 0.001 |
| postcentral gyrus (S1) | 8 | R | 56 | -20 | 50 | 5.05 | 6.12 | 0.004 |
| cerebellum | 1 | L | -11 | -65 | -54 | 4.96 | 5.96 | 0.026 |
| midcingulate cortex | 9 | R | 12 | -25 | 38 | 4.89 | 5.84 | 0.003 |
| postcentral gyrus (S1) | 8 | L | -31 | -35 | 70 | 4.88 | 5.83 | 0.004 |
| rolandic operculum | 3 | R | 52 | -5 | 8 | 4.84 | 5.76 | 0.013 |
| occipital cortex | 1 | L | -16 | -78 | 23 | 4.82 | 5.74 | 0.026 |
| task<br>**FGAT-distant** | | | parametric modulator<br>**response time during the FGAT-distant** | | | | | |
| cluster label | cluster size | peak side | x | y | z | Z-score | T-value | cluster p-value |
| **supramarginal gyrus/superior temporal gyrus** | **237** | **R** | **62** | **-48** | **28** | **5.92** | **7.71** | **5.10-11** |
| **lateral frontal pole** | **216** | **R** | **32** | **50** | **23** | **5.86** | **7.59** | **2.10-10** |
| superior frontal gyrus/middle frontal gyrus | 56 | L | -28 | -92 | 23 | 5.52 | 6.94 | 1.10-4 |
| occipital cortex | 21 | L | -28 | 50 | 13 | 5.45 | 6.82 | 0.009 |
| superior frontal gyrus/middle frontal gyrus | 13 | R | 36 | 20 | 8 | 5.34 | 6.62 | 0.033 |
| insula/inferior frontal gyrus | 14 | L | -44 | -52 | -47 | 5.15 | 6.29 | 0.028 |
| cerebellum | 14 | L | -36 | 38 | 40 | 5.13 | 6.25 | 0.028 |
| fuisform gyrus/lingual gyrus | 16 | R | 24 | -68 | -14 | 5.10 | 6.20 | 0.020 |

**Supplementary Table 2: All clusters in the SPMs of the parametric modulation analyses of the likeability rating task and the FGAT-distant by response times.** p-values are FWE corrected. Coordinates x, y and z refer to the Montreal Neurological Institute space. Only the results in bold are reported in the supplementary results.

### A NETWORK TIMESERIES APPROACH

At the behavioral level, originality and adequacy contribute to likeability ratings : $L = (\alpha O^{\delta} + (1\text{-}\alpha)A^{\delta})^{1/\delta}$

Do DMN and ECN activities contribute similarly to the BVS activity: $BVS = (\alpha DMN^{\delta} + (1\text{-}\alpha)ECN^{\delta})^{1/\delta}$ ?

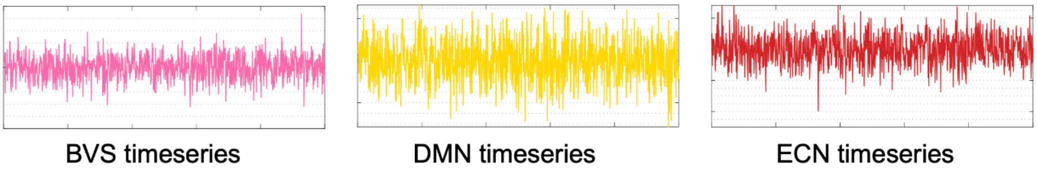


### B CORRELATION BETWEEN BEHAVIORAL PARAMETERS AND NEURAL PARAMETERS ESTIMATED WITH THE *FGAT-DISTANT* TIMESERIES

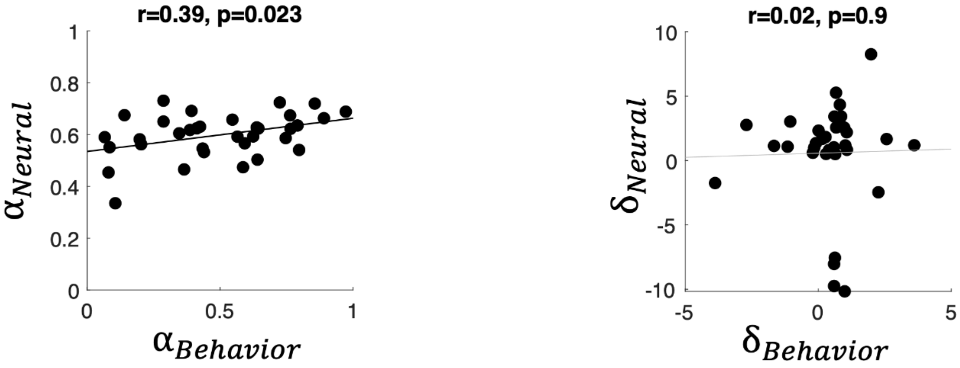


### C CORRELATION BETWEEN BEHAVIORAL PARAMETERS AND NEURAL PARAMETERS ESTIMATED WITH THE *RESTING-STATE* TIMESERIES

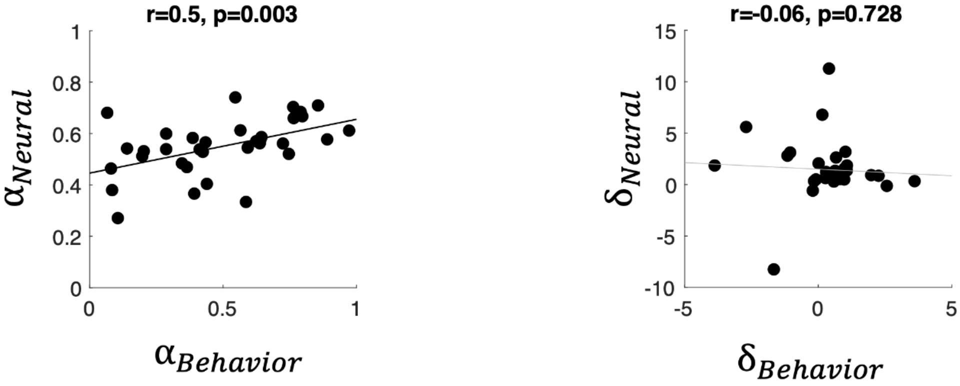

**Figure S3: Correlation between the parameters of the CES, estimated with behavioral and neural data. (A)** Schematics of the analysis approach. The timeseries depicted here for illustration purposes are simulated data. **(B)** Correlation between the behavioral parameters and the neural parameters estimated using the FGAT-distant timeseries. **(C)** Correlation between the behavioral parameters and the neural parameters estimated using the resting-state timeseries. $\alpha_{behavioral}$ and $\delta_{behavioral}$ were estimated using the CES model fitted on the likeability, originality and adequacy ratings. $\alpha_{neural}$ and $\delta_{neural}$ were estimated using the CES model fitted on BVS, DMN and ECN timeseries during the FGAT-distant task (B) or during the resting-state (C). Each dot represents one participant. Black lines indicate significant correlations ($p<0.05$), and grey lines indicate non-significant correlations ($p>0.05$).
